# Crude *Fucus vesiculosus* fucoidan demonstrates superior SARS-CoV-2 antiviral activity compared to its pure form: binding kinetics and functional studies

**DOI:** 10.64898/2026.05.07.723385

**Authors:** Anna Dudek, Rajendra Prasad Janapatla, Chyi-Liang Chen, Cheng-Hsun Chiu

**Author notes:** Correspondence: Cheng-Hsun Chiu, Division of Pediatric Infectious Diseases, Department of Pediatrics, Chang Gung Memorial Hospital, Chang Gung University College of Medicine, Taoyuan, Taiwan. Equal contribution: Anna Dudek, Rajendra Prasad Janapatla.

## Abstract

Fucoidans have been widely reported to show SARS-CoV-2 antiviral activity. In this study, we observed a striking difference in the inhibitory potency between two commercially available fucoidans: *Fucus vesiculosus* crude (Fvc) and pure (Fvp). SEC-MALS analysis revealed two molecular weight populations for Fvc (1098 kDa, 58.58 kDa) and one for Fvp (40.48 kDa). At micromolar concentrations of fucoidans, the binding affinities (KDs) of Fvc_1098 (223 nM) and Fvc_58 (4.27 µM) for the amine-biotinylated SARS-CoV-2 receptor binding domain (RBD) were higher than that of Fvp (76.5 µM). At nanomolar concentrations, binding was observed only to the Avi-tag-, but not amine-biotinylated RBDs, suggesting better accessibility of their binding sites. The association rates (k_on_) were faster for Fvc than for Fvp. Similarly, affinities of Fvc_1098 (23.4 nM) and Fvc_58 (4.48 μM) for ACE2 were greater than that of Fvp (66.8 μM), indicating that Fvc can bind directly to both RBD and ACE2. Fvc demonstrated enhanced inhibitory potency (IC50 = 58 μg/mL) compared to Fvp (IC50 > 239 μg/mL) in the pseudovirus entry assay and did not induce cytotoxicity in HEK293T cells. In conclusion, crude fucoidan with high fucose content and high molecular weight shows promising antiviral activity.

## 1. Introduction

Severe acute respiratory syndrome coronavirus 2 (SARS-CoV-2) usually causes respiratory complications, but it can also affect the gastrointestinal tract [1, 2]. The receptor-binding domain (RBD) of the S1 subunit of the spike protein specifically interacts with the angiotensin-converting enzyme 2 (ACE2) receptor on host cells, enabling viral attachment and subsequent infection [3]. Proceeding ACE2 interaction, RBD associates with glycosaminoglycan (GAG) heparan sulfate, a co-receptor on host cells, through its GAG-binding motif (YRLFRKS) [4–6]. It has been reported that the ancestral SARS-CoV-2 spike RBD has a binding affinity for the ACE2 receptor in the nanomolar range [7–9]. Mutations in the spike proteins of the Alpha, Beta, Delta, and Omicron variants have been found to enhance their binding affinities for the ACE2 receptor [10, 11]. Recently, researchers from the Wuhan Institute have identified a new bat merbecovirus, HKU5-CoV-2 lineage 2, capable of infecting human ACE2-expressing cells, raising concerns about its potential transmission to humans [12]. Therefore, studying the interactions between the spike protein and their receptors on host cells, as well as identifying inhibitors to block these processes, is crucial for the development of therapeutic agents. Especially since the continued emergence of new variants/mutants has caused a reduction in the efficacy of vaccine candidates and monoclonal antibodies targeting the spike protein [13–15].

Many reports have indicated that the binding affinities of sulfated polysaccharides for the spike protein correlate with their *in vitro* antiviral activities. Sulfated polysaccharides, such as heparan sulfate, heparin, and seaweed sulfated fucose polymers (fucoidans), have demonstrated binding affinities for the spike protein, mostly in the micro- to nanomolar range, depending on factors such as molecular weight (MW), degree of sulfation, and its pattern [5, 16–19]. Fucoidans and other seaweed polysaccharides are generally recognized as safe (GRAS), and have been shown to inhibit the initial steps of viral attachment and fusion to host cells by mimicking GAG chemical structure [5, 16, 20], blocking spike protein GAG binding motifs and cellular receptors [21, 22], thereby preventing infection [23]. Fucoidans have been classified into three types based on the linkage of their L-fucopyranosyl residues: α(1→3) (Type I), alternating α(1→3) and α(1→4) (Type II), and α(1→4)-linked (Type III) [17, 24]. The backbone of *Fucus vesiculosus* fucoidan (Fv) has been found to be Type II with the sulfation pattern typically at the C-2 and C-4 but also at the C-3 positions of the fucopyranosyl residues [17, 24–26]. Detailed analysis of the fucose glycosidic linkages of Fv revealed 1,3,4-linked (26.5 %), followed by 1- and 3-linked (24.2 %), 1- and 4-linked (14.2 %), and 1,2-linked (6.9 %), with a sulphate group at position 2, found at branching units, with glycosidic bonds at positions 1, 3, and 4 [26].

Clinical trials have confirmed that oral fucoidans (1 g/day) from *F. vesiculosus* and *Undaria pinnatifida* are well-tolerated [27], with no adverse events reported [28]. Although SARS-CoV-2 spreads mainly through respiratory droplets, detection of viral RNA in fecal samples raises concerns about potential fecal-oral transmission [29–31]. The expression of ACE2 throughout the gastrointestinal surface suggests, that it could serve as an alternative site for SARS-CoV-2 entry and infection [32, 33]. Fucoidans show promise as candidates for further investigation as oral prophylactic and therapeutic agents for SARS-CoV-2 and other ACE2-dependent viral infections, particularly in preventing viral entry into gastrointestinal tract cells. Considering the potential of marine polysaccharides to inhibit RBD binding to ACE2, the aim of this study was to perform an initial ELISA-based screening of commercially available marine polysaccharides such as fucoidans and alginates to assess their binding and inhibitory potency against RBDs from different SARS-CoV-2 variants. The results showed that the most potent inhibitor was crude fucoidan from *F. vesiculosus* (Fvc) while its pure form (Fvp) remained inactive. Fvc (Cat. no. F5631) and Fvp (Cat. no. F8190) are commercially available products from Sigma-Aldrich. According to the manufacturer, information on the purification process of Fvp and its relationship to Fvc is not available. We hypothesize that the crude fucoidan Fvc exhibits higher binding affinities for RBD and ACE2 than its pure form Fvp due to a mechanism facilitated by its high molecular weight. This hypothesis was supported by molecular weight analysis using SEC-MALS, binding kinetics measured by biolayer interferometry, and inhibitory effects evaluated using pseudovirus entry assay. The results showed that the crude fucoidan with high molecular weight demonstrates promising antiviral activity.

## 2. Materials and methods

### 2.1. Materials

Fucoidans from *Fucus vesiculosus* crude (Fvc) (Cat. No. F5631, Lot No. SLCD5199) and pure (Fvp) (Cat. No. F8190, Lot No. SLCB9091, purity ≥ 95 %) were purchased from Sigma-Aldrich (St. Louis, MO, USA). SARS-CoV-2 receptor-binding domain (RBD) proteins (Wuhan-Hu-1, Alpha B.1.1.7, Beta B.1.351, Delta B.1.617.2, and Omicron BA.4&BA.5), angiotensin-converting enzyme 2 (ACE2), ACE2 derived spike-binding peptide 1 (SBP1), and other polysaccharides were purchased from commercial sources listed in the Supplementary Table 1 and 2.

### 2.2. Chemical composition of fucoidans

The monosaccharide composition of fucoidan samples was analyzed using High Performance Anion-Exchange Chromatography with Pulsed Amperometric Detection (HPAEC-PAD) (Dionex ICS-3000 system, Thermo Scientific) equipped with a CarboPac PA20 analytical column (3 × 150 mm) and a corresponding guard column. The injection volume was 10 µL with a flow rate of 0.25 mL/min. The column temperature was maintained at 20 °C. The detection was performed using pulsed amperometric detection with the waveform setting “Gold, Carbo, Quad” and an Ag/AgCl reference electrode. Fucoidans (1 mg/100 µL) were treated with 2 M trifluoroacetic acid (TFA) at 99 °C for 4 h, SpeedVac dried, dissolved in 100 µL of deionized water, diluted 10-fold, and 0.22 µm filtered. The injection volume was set to 10 μL. Neutral monosaccharide standards (fucose (Fuc), galactose (Gal), glucose (Glc), mannose (Man), xylose (Xyl)) and uronic acid standards (galacturonic acid (GalA), guluronic acid (GulA), glucuronic acid (GlcA), mannuronic acid (ManA)) were used at concentrations of 2 µM and 10 µM each. The spike-in concentrations of GalA and ManA were 2 µM (20 pmol/10 µL) each. For neutral monosaccharides, the isocratic elution was performed with 14 mM NaOH from 0 to 17 min, followed by a 5-min high-strength wash with 200 mM NaOH (17-22 min). For acidic monosaccharides, a gradient of sodium acetate in 100 mM NaOH was applied as follows: 0–20 min, 76-200 mM NaOAc; 20-25 min, 200 mM NaOAc; 25-30 min, followed by a 5-min high-strength wash with 500 mM NaOAc. The monosaccharide concentrations were determined using the standard curves of the peak areas plotted against the known standard concentrations.

The sulfate analysis was performed using a modified sulfate assay kit (KA1621, Abnova, Taiwan). The method utilizes the quantitative formation of insoluble barium sulfate in polyethylene glycol. Fucoidan samples were incubated in 0.2 M HCl at 80 °C overnight to liberate sulfate ions. Absorbance was measured at 570 nm. A serially diluted sulfate standard was used to generate a calibration curve (Supplementary Fig. 1A).

The total uronic acid content was measured using a colorimetric assay [34]. A volume of 200 μL of each fucoidan sample and a serially diluted alginic acid standard (alginic acid sodium salt from brown algae (Sigma-Aldrich, USA)) was added to 1.2 mL of 0.0125 M tetraborate solution (in concentrated sulfuric acid), vortexed, placed on ice for 10 min, incubated at 100 °C for 5 min, and put back on ice for another 5 min. Next, 20 μL of 0.15% (w/v) m-hydroxybiphenyl in 0.125 M NaOH (Sigma Aldrich, USA) was added and samples were vortexed for 1 min. Absorbance was measured at 520 nm, and the blanks (20 μL of m-hydroxybiphenyl was replaced by 20 μL of 0.125 M NaOH) were subtracted from both samples and standards. Uronic acid concentration was calculated based on the standard curve for alginic acid (Supplementary Fig. 1B).

The total carbohydrate content was measured by microplate-based phenol-sulfuric acid (PSA) method proposed by Ogura et al. [35]. Fucoidans (Fvc, Fvp) and a serially diluted glucose standard (5 μL of 10 mg/mL in 25 μL of water) in 150 μL of sulfuric acid were heated at 90 °C for 15 min. Then, a volume of 30 μL of 5% (w/v) phenol was added, the plate was shaken for 5 min, and allowed to rest for over 60 min. Absorbance was measured at 490 nm (Supplementary Fig. 1C).

### 2.3. Size-exclusion chromatography with multi-angle light scattering (SEC-MALS)

The molecular weights of Fvc and Fvp were determined using a multi-angle laser light scattering (MALS) system, which included a multi-angle laser light-scattering photometer (DAWN HELEOS II; Wyatt Technology, Santa Barbara, CA, USA), a differential refractive index detector (Optilab T-Rex; Wyatt Technology, USA), and a UV-absorbance detector. The system operated with 1x PBS (phosphate-buffered saline) containing 200 ppm sodium azide at a flow rate of 0.5 mL/min. A 100 µL sample was injected into a WTC-30S5 7.8 × 300 mm column (Wyatt Technology, USA), and data collection and analysis were performed using ASTRA software (Wyatt Technology, USA). The differential refractive index increment (dn/dc) was determined using the saccharide method, with a value of 0.147 for saccharide.

### 2.4. Staining of fucoidans in polyacrylamide gel

Fucoidans (100 µg/mL in 62.5mM Tris-HCl buffer, pH 7.4, containing 10 % glycerol and 0.01% phenol red) were electrophoresed on a gradient gel (Acrylamide WB F1 Gradient Gel 4-20%, Future Scientific Co., LTD, BioFuture, Taiwan) in 1 x 25 mM Tris, 192 mM glycine, and 0.1 % (w/v) SDS running buffer, pH 8.3 (Bio-Rad Laboratories, Inc., USA). Gels were stained with 0.01 % *O*-toluidine blue in 2 % acetic acid for 1-2 h and destained with 2 % acetic acid (2 x 30 min), followed by water until the desired bands were observed. Gels were also stained using the Thermo Scientific Pierce Silver Stain Kit and Bio-Rad Silver Stain according to the manufacturer’s modified protocol optimized for minigels. The sequential steps of both kits include fixation, sensitization, silver impregnation/enhancement, and development. In the Bio-Rad kit, potassium dichromate oxidizes hydroxyl groups of fucoidan, enabling silver ion binding. The chemical composition of the Thermo Scientific Pierce Silver Stain Kit reagents is not provided by the manufacturer.

### 2.5. Biolayer Interferometry (BLI)

To evaluate the binding kinetic parameters, including the association rate constant (k_on_) and equilibrium dissociation constant/binding affinity (KD), for the interactions of Fvc and Fvp with RBDs, ACE2, and SBP1, BLI experiments were conducted using Octet RED96 instrument (ForteBio, Fremont, CA, USA). Amine- (25 µg/mL) and Avi-tag-biotinylated RBDs (12.5 to 6.25 µg/mL), biotinylated ACE2 (12.5 µg/mL), in PBSTB (1 x PBS (Bioman Scientific CO, LTD., Taiwan) + 0.02 % Tween 20 + 0.1 % bovine serum albumin (BSA)) were loaded on streptavidin (SA) biosensors pre-incubated in PBSTB for 30 min. Fucoidans (5000 to 5 µg/mL) serially diluted in PBSTB were used as analytes. The Fvc analyte, at the highest tested concentration of 5000 µg/mL in PBSTB, did not bind to the unloaded SA biosensor, resulting in no observable increase in the BLI response signal (RS). Biotinylated SBP1 (25 µg/mL) was loaded onto high-precision streptavidin (SAX) or SA biosensors in PBS buffer to assess its affinity for the Omicron BA.4&BA.5 RBD or Fvc and Fvp, respectively. The association and dissociation phases were monitored for 600 s. Data were reference-subtracted (ligand-loaded SA or SAX biosensors in PBSTB or PBS), and binding kinetic parameters were derived using the Octet Data Analysis Software v10.0 (ForteBio, USA), applying a 1:1 binding affinity model for protein and peptide ligands in either global or local fit mode. Since SEC-MALS analysis of Fvc revealed two populations with molecular weights of 1098 kDa and 58.58 kDa, separate KD values were calculated for each molecular weight assuming the 1:1 binding model. Experiments were performed in at least duplicate using new biosensors for each experiment and analyte concentration.

### 2.6. Effect of polysaccharides on pseudovirus infection of HEK293T-ACE2-OE cells

ACE2-expressing HEK293T cells (HEK293T-ACE2-OE) were seeded at 0.75 × 10^5^ cells per well in 24-well plates. After 48 h, confluent HEK293T-ACE2 cells were infected with 5 µL of 10^7^ TU/mL lentiviral particles pseudotyped with SARS-CoV-2 and carrying GFP as a reporter gene (ACE Biolabs, Taiwan). For the inhibition assay, various concentrations of Fvc, Fvp, alginate from *Ascophyllum nodosum* (AlgAn), and mannan (Man) were pre-incubated with pseudotyped virus for 2 h. The mixture was then transferred to the confluent HEK293T-ACE2 cells and incubated for 24 h to allow for infection. Following the 24-h infection period, the medium was replaced, and the number of GFP-positive cells was quantified between 48- and 72-h post-infection using a Cytation™ 3 Cell Imaging Multi-Mode Reader (BioTek Instruments, Winooski, VT, USA).

To analyze the cytotoxicity of polysaccharides, HEK293T-ACE2-OE cells (1 × 10^4^ cells/well) were seeded in 96-well plates and incubated at 37 °C in a 5 % CO₂ incubator for 48 h [36]. The confluent monolayers were then treated with various concentrations of polysaccharides for 24 h. After incubation, 10 µL of WST-1 reagent was added to each well, and the cells were incubated for an additional 2-4 h. Absorbance was measured at 450 nm using a microplate reader (SpectraMax M5 plate reader, Molecular Devices, USA). Cell viability was calculated as a percentage relative to the untreated control group.

### 2.7. Statistical analysis

Data (presented as mean ± SEM) were analyzed using the Shapiro-Wilk normality test and a two-way repeated-measures analysis of variance (ANOVA), followed by Bonferroni’s multiple comparisons test (GraphPad Prism version 10.4.1 for Windows, GraphPad Software, San Diego, CA, USA). Results with p < 0.05 were considered statistically significant.

## 3. Results and discussion

### 3.1. Reactivity of marine polysaccharides toward RBDs

The interaction of RBD proteins with various polysaccharides was assessed using Carbo-BIND ELISA microplates (Supplementary Materials and methods 1.1). All tested RBD variants demonstrated the strongest binding to the coated Fvc (OD > 3.0) (Supplementary Fig. 2A-B). Moderate reactivity (OD < 2.0) was observed with other sulfated polysaccharides, including fucoidan from *A. nodosum*, heparin, Κ-carrageenan, dextran, and other commercially available fucoidans, as well as with non-sulfated polysaccharides such as algi nates. Differences in the binding potency of tested polysaccharides could be due to several factors, including molecular weight and the structure of fucoidans, which can vary significantly in branching, monomer residue linkages, sulfation, and acetylation depending on the algae species and harvesting season [37]. In the ELISA-based inhibition assay, Fvc (IC50 = 26 μg/mL), Fvc nutritional grade from Marinova (IC50 = 53 μg/mL), and alginate from *A. nodosum* (IC50 = 61 μg/mL) were the most effective inhibitors among the polysaccharides tested (Supplementary Fig. 2C). Of note, Fvp failed to inhibit the RBD activity. Song et al. [18] reported similar IC50 values for the inhibition of spike glycoprotein binding to a heparin-coated SPR chip: 65.6 μg/mL for high molecular weight fucoidan from *Saccharina japonica*, and 53.9 μg/mL for rhamnan sulfate from *Monostroma nitidum*. In contrast, Koike et al. [17] found that a low molecular weight fucoidan derivative exhibited the highest inhibitory activity among the fucoidan derivatives tested (IC50 = 86.9 μM). Based on the ELISA results, Fvc and Fvp were selected for further characterization.

### 3.2. Characterization of Fvc and Fvp

Monosaccharide composition analysis of Fvc revealed that it predominantly consists of fucose (87.41 %), with trace amounts of Gal, Glc, Man, Xyl, GlcA, and ManA (0.13 - 4.63 %) (Table 1, Supplementary Fig. 3). This aligns with previous reports for Fvc, showing 86.2 % fucose with minor monosaccharides (2 - 5.1 %) by GC-MS [38], and 82.6 % fucose (0.4 - 6.8 %, including GlcA and ManA) by HPAEC-PAD [39]. The observed difference in monosaccharide composition may be attributed to variations in analytical methodologies and the use of different lots of Sigma-Aldrich fucoidan samples. Fucose content in Fvp (60.15 %) was lower than that in Fvc, however other monosaccharides were present at significantly higher levels (0.21 - 13.75 %) (Table 1). After optimizing the spike-in concentrations of GalA and ManA standards, the ManA peaks at 22.1 min (Fvc) and 21.9 min (Fvp) were confirmed, however, the minor peaks at 18.7 min present in Fvc and Fvp samples, could not be identified as GalA (Supplementary Fig. 4). The lower fucose content and higher neutral and acidic monosaccharide impurities in Fvp may result from purification that causes depolymerization and loss of fucose components, potentially reducing its bioactivity. Fvc showed a higher fucose level due to a greater abundance of intact fucose polymers. The sulfate content of both Fvc and Fvp was similar, however total uronic acid and total carbohydrate content were higher for Fvc (28.3 %, 7.2 %, 63.6 %) than for Fvp (28.9 %, 5.1 %, 59.7 %) (Table 1), suggesting that the purification process preserves sulfation level and partially removes uronic acid contaminants and other carbohydrate components.

**Table 1.** Chemical composition of Fvc and Fvp.

|  | Monosaccharides [mol %]<sup>a</sup> |  |  |  |  |  |  | Uronic acid [%]<sup>b</sup> | Total carbohydrate [%]<sup>c</sup> | Sulfate [%] |
| --- | --- | --- | --- | --- | --- | --- | --- | --- | --- | --- |
|  | Fuc | Gal | Glc | Man | Xyl | GlcA | ManA |  |  |  |
| Fvc | 87.41 | 4.63 | 0.63 | 0.99 | 3.54 | 2.67 | 0.13 | 7.2 ± 0.1 | 63.6 ± 0.5 | 28.3 ± 0.8 |
| Fvp | 60.15 | 11.87 | 13.75 | 1.39 | 7.65 | 4.98 | 0.21 | 5.1 ± 0.1 | 59.7 ± 0.7 | 28.9 ± 0.6 |

<sup>a</sup> HPAEC-PAD; <sup>b</sup> colorimetric assay; <sup>c</sup> PSA method

The SEC-MALS analysis of Fvc revealed a wide, asymmetric, bimodal peak indicating a high degree of molecular weight heterogeneity [40]. Two peaks were observed: a narrow, more intensive Peak1 with a maximum at 10.79 min and a normalized light scattering (LS) intensity of 0.915, and a broad, less intense Peak2 with a maximum at 13.97 min, and a normalized LS intensity of 0.477, suggesting the presence of two different molecular weight populations (Fig. 1A). The initial intense Peak1 revealed high molecular weight chains with an average MW of 1098 ± 4.749 kDa (range: 500 to 15,000 kDa, Fvc_1098), while the second, less intensive Peak2 corresponded to a lower molecular weight population with an average MW of 58.58 ± 4.199 kDa (range: 32 to 120 kDa, Fvc_58). In contrast, SEC-MALS analysis of Fvp yielded a single peak with its maximum at 12.5 min and a normalized LS intensity of 0.926, indicating a lower average MW of 40.48 ± 1.38 kDa (range: 10 to 295 kDa, Fvp) (Fig. 1A).

**Fig. 1.**
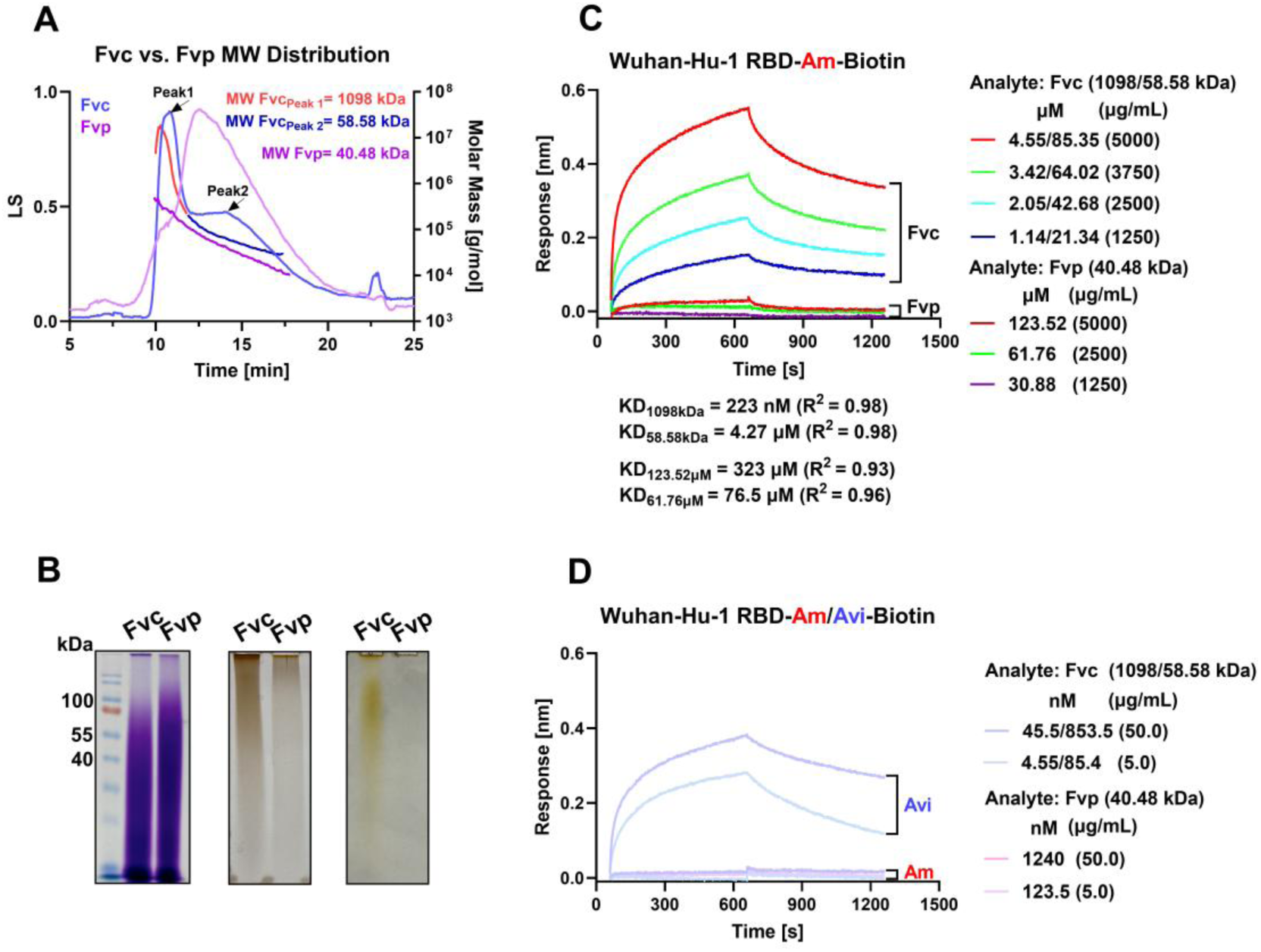
Crude (Fvc) and pure (Fvp) fucoidan characteristics. SEC-MALS chromatograms of Fvc (blue line) and Fvp (purple line) along with their molecular weight trace lines: Fvc Peak1 (red line), Fvc Peak2 (navy blue line), and Fvp (purple line) (A), Electropherograms of Fvc and Fvp stained with *O*-toluidine blue (left), Thermo Scientific Pierce Silver Stain Kit (middle), Bio-Rad Silver Stain (right) (B), Sensorgrams of Fvc and Fvp binding to the amine- (Am) and Avi-tag-(Avi) biotinylated Wuhan-Hu-1 RBD ligand (C) and (D).

Fvc and Fvp appeared as smeared bands upon *O*-toluidine blue staining. Silver staining also revealed a smeared band mainly in the higher molecular weight region for Fvc, with no such band observed for Fvp (Fig. 1B). This is consistent with the SEC-MALS results, which detected high molecular weight chains only in Fvc but not in Fvp. Polysaccharides containing oxidizable groups and negatively charged sulfate groups can bind positively charged silver which is then reduced to metallic silver producing a dark stain [41]. Fvp was not effectively stained, which may be caused by the alteration or decreased reactivity of functional groups capable of binding silver after the purification process.

The proteomic profiles of Fvc and Fvp solutions and gels were analyzed using the marine algal databases (Supplementary Materials and methods 1.2). More proteins were identified in the Fvc solution compared to the Fvp solution, though only a few unique peptides (ranging from 1 to 4) were observed, suggesting that these trace proteins may be contaminants. No substantial differences were observed between Fvc and Fvp in the representative excised gel regions of ∼100 kDa (Supplementary file 2). A recent study using LC-MS reported the presence of 77 metabolites in Fvc and only 35 in Fvp (both sourced from Sigma-Aldrich) [42]. These differences may contribute to the observed variations in the ELISA binding and inhibitory potency between Fvc and Fvp. Further research is needed to identify the active metabolites and to rule out their potential interference in fucoidan inhibitory activity.

Figure 1C shows BLI sensorgrams depicting the association and dissociation profiles of Fvc and Fvp with the RBD, biotinylated via amines (Wuhan-Hu-1 RBD-Am-Biotin), at micromolar concentrations of fucoidans. The KD values of high and low molecular weight Fvc_1098 and Fvc_58 for Wuhan-Hu-1 RBD-Am-Biotin were determined to be 223 nM and 4.27 µM respectively, within the expected micromolar range for moderate affinity carbohydrate-protein interactions [43, 44]. Liu et al. [45] found the affinity of heparin for the RBD was also in micromolar range (1 μM) using SPR as a method.

The Fvc analyte displayed a response signal (RS = 0.55 nm) that was nearly 20-fold higher than that of Fvp (RS = 0.029 nm) at the highest concentration tested (5000 µg/mL), probably due to the presence of high molecular weight fraction in Fvc. Correspondingly, the binding affinity of Fvp for the Wuhan-Hu-1 RBD-Am-Biotin was lower than that of Fvc_1098 and Fvc_58, with a KD range of 76.5 to 323 µM. Purification conditions may lead to depolymerization of fucoidans, and the loss of other carbohydrates and bioactive components [40].

When the concentrations of the Fvc and Fvp analytes were diluted 100-fold to 50 µg/mL, no binding was observed to Wuhan-Hu-1 RBD-Am-Biotin (Fig. 1D). To test the hypothesis that the lack of binding was due to the biotinylation of the RBD protein via amine groups, which could potentially block carbohydrate-binding sites, Avi-tag-biotinylated RBD was used [46]. In this Avi-tagged protein, a single lysine residue in the Avi-tag peptide was enzymatically labeled with biotin, allowing for uniform protein orientation and bioactivity preservation as the rest of the protein remained unmodified, as described on the manufacturer’s website [47]. As shown in the sensorgrams in Fig. 1D, interactions between Wuhan-Hu-1 RBD-Avi-Biotin and Fvc generated a significantly higher BLI response signal (RS = 0.32 nm) at the analyte concentration of 50 µg/mL compared to the response observed with Wuhan-Hu-1 RBD-Am-Biotin and Fvc (RS ≤ 0.015 nm). The association rate constants of Fvc_1098 and Fvc_58 ranged from 2.47 × 10^4^ to 1.21 × 10^6^ M⁻¹s⁻¹, with corresponding KD values in the nanomolar range (1.33 to 28.1 nM), indicating high binding affinity. Similarly, Xu et al. [19] reported that a high molecular weight sulfated exopolysaccharide from *Haloarcula hispanica* bound to the RBD with high affinity (KD = 2.23 nM), blocking spike protein binding to Vero E6 and bronchial epithelial BEAS-2B cells and inhibiting pseudovirus infection.

### 3.3. Association rates of Fvc and Fvp with Avi-tag-biotinylated RBDs

The binding kinetics of Fvc and Fvp with Avi-tag-biotinylated RBDs from different SARS-CoV-2 variants including Alpha, Beta, Delta, and Omicron were analyzed to determine the k_on_ and KD values. The BLI response signals generated from the interaction of RBD variants, such as Alpha and Beta, with Fvc were higher (0.19 to 1.6 nm) compared to those with Fvp (< 0.45 nm) at higher fucoidan concentrations (46.3 to 1250 μg/mL), suggesting the binding of high molecular weight Fvc components or their aggregates (Supplementary Fig. 5A-B). However, at lower concentrations (5 and 50 μg/mL), the BLI response signals for Fvc were similar to those of Fvp (Supplementary Fig. 5E-F) and they increased progressively across the Alpha, Beta, Delta, and Omicron RBDs (averaged RS values for Fvp and Fvc: 0.27, 0.57, 0.76, and 0.81 nm, respectively), correlating with the increasing number of mutations (1, 3, 2, and 17, respectively) and corresponding increases in net positive charge (6.5, 7.5, 8.5, 10.6; Supplementary Fig. 6). The increased RS values could be attributed to enhanced electrostatic interactions between the negatively charged fucoidan and the positively charged residues within the RBD, including the conserved GAG-binding motif [18]. Kwon et al. demonstrated that suramin—a polysulfated polyanion—binds more tightly to the Omicron RBD than to the wild type or Delta variants due to key electropositive mutations (e.g., N440K, T478K, Q493R, Q498R, Y505H) that alter RBD electrostatic surface and introduce new sulfate-interacting residues (e.g., Arg493). SPR and molecular docking data showed that suramin inhibits the binding of the Omicron RBD to both ACE2 and heparan sulfate receptors [36]. These findings are consistent with our observation that variants with a higher net positive charge show stronger fucoidan binding. In addition, other structural features may also influence antiviral activity, as the negatively charged chondroitin sulfate demonstrated minimal inhibitory potency [48].

Next, the k_on_ values were determined for all the Fvc-RBDs and Fvp-RBDs interactions, providing further insight into the binding kinetics. For the Avi-tag-biotinylated RBD ligands, Fvc_1098 consistently demonstrated increasing k_on_ values as the concentration decreased. The calculated k_on_ values of Fvc_1098 ranged from 5.02 × 10^3^ to 1.62 × 10^5^ M^-1^s^-1^across the concentration range of 1250 to 46.3 μg/mL (Fig. 2A). This trend suggests that rate of association increases at lower concentrations. In contrast, Fvc_58 and Fvp consistently exhibited lower k_on_ values across all concentrations compared to Fvc_1098. The k_on_ values of Fvc_58 and Fvp with RBDs (Alpha, Beta, Delta, and Omicron) ranged from 6.27 × 10^1^ to 1.56 × 10^4^ M^-1^s^-1^ (Fig. 2B-C), suggesting that Fvc_58 and Fvp generally exhibit slower or less efficient association kinetics, which may result in higher KD values [46, 49]. This observation is consistent with the findings of Chang et al. [49], where anti-SARS-CoV-2 antibodies with the highest KD values displayed lower on-rates, indicating that slower association can contribute to reduced binding efficiency. The faster association rate of Fvc_1098, compared to Fvc_58 and Fvp, may be attributed to multivalent interactions between the high molecular weight Fvc and the RBDs. At lower concentrations (≤ 50 μg/mL), the k_on_ values of Fvc_1098 (2.07 × 10^5^ to 5.49 × 10^6^ M^-1^s^-1^) were comparable to those of potent monoclonal antibodies targeting the RBD or S1 domain of the spike protein [50]. In contrast, the k_on_ values reported for interactions between carbohydrate-binding proteins and glycoconjugates or polysaccharides were found to range from 10^2^ to 10^5^ M^−1^s^−1^ [44, 51]. The KD values for Fvc and Fvp, used in nanomolar concentrations, with Avi-tag-biotinylated RBD proteins of the Alpha, and Delta variants, were in the nanomolar range. However, the KD values for Fvc and Fvp with Beta and Omicron RBDs were below the picomolar threshold (< 1 × 10^-12^ M), reaching the limit of detection of the BLI assay (Supplementary Fig. 5E-F, and 5I-J). While this suggests particularly strong binding, caution is warranted due to the instability of the Avi-tag biotinylated ligands (Supplementary Fig. 5G-H, and 5K-L). This instability hindered the acquisition of reliable association-dissociation curves and prevented accurate calculation of KD values. Nevertheless, an increase in the response signal between the reference (RBD-loaded SA biosensor in PBSTB only) and the sample (RBD-loaded SA biosensor in fucoidan dissolved in PBSTB) still indicated binding of fucoidans to the RBDs. This limitation could be partly addressed by using biotinylated Fvc as the ligand, which confirmed the binding of Omicron RBD to fucoidan; however, the response signal was low (0.08 nm) (Supplementary Fig. 7). This may be due to the random biotinylation of fucoidan following its amination, which could potentially interfere with its binding ability. The above results indicate that mutations in the RBDs had minimal effect on the association rates of Fvc and Fvp at nanomolar concentrations, consistent with the observation using Avi-tag-biotinylated Wuhan-Hu-1 RBD protein. Instead, the method of RBD biotinylation played a more critical role. Biotinylation through amine groups may obstruct binding sites targeted by Fvc and Fvp, whereas terminal biotinylation preserves the accessibility of these sites [46]. He et al. [23] found that the binding affinities of the RBD for heparin were in the nanomolar range, with the XBB.1.5 variant (KD = 160 nM) exhibiting slightly stronger affinity than the ancestral strain (KD = 350 nM), as determined by SPR.

**Fig. 2.**
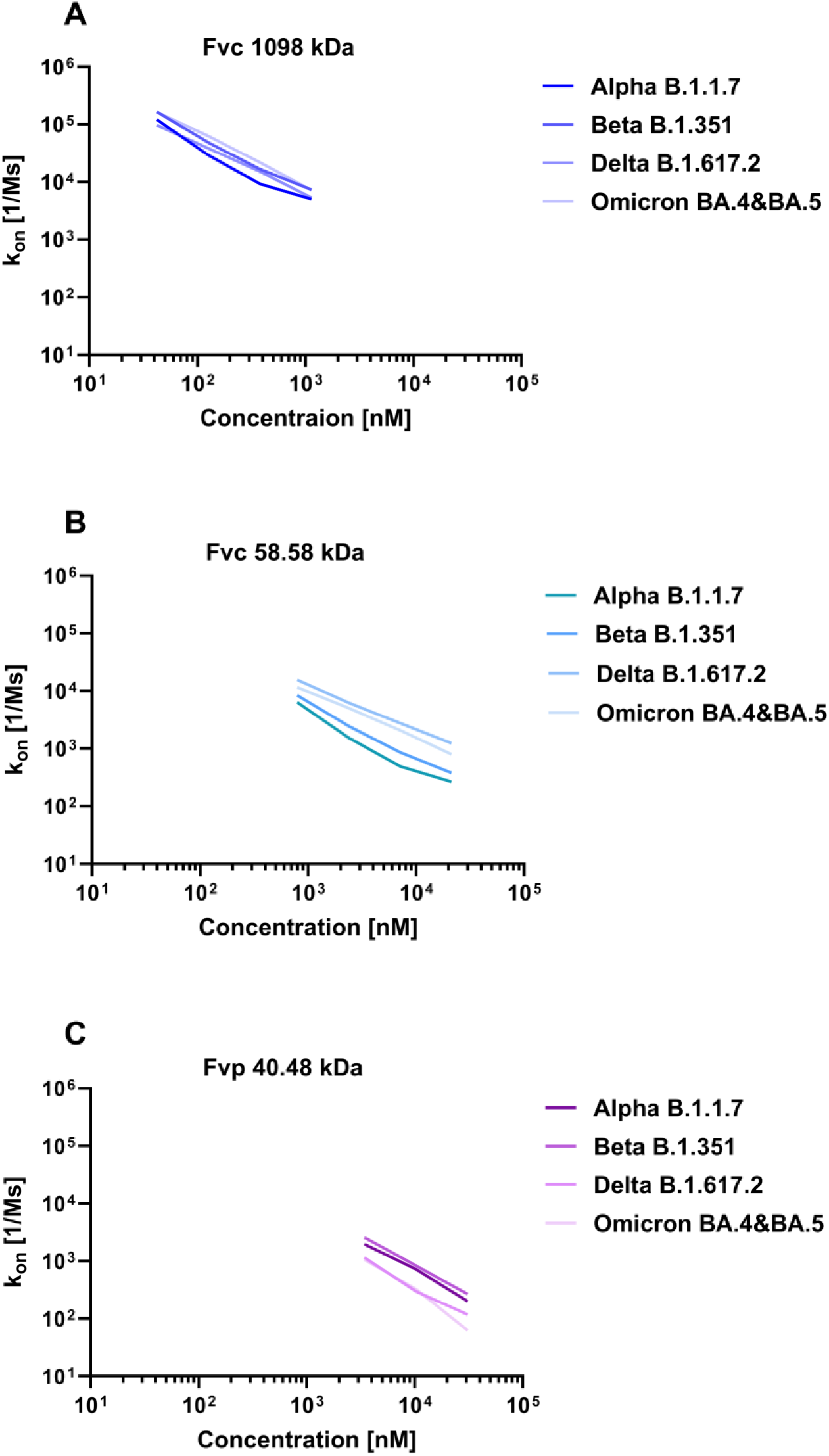
Association rate constants (k_on_) of Fvc_1098 (A), Fvc_58 (B), and Fvp (C) against their concentrations, showing how quickly fucoidans bind to RBDs.

### 3.4. Binding affinities of Fvc and Fvp for ACE2 and SBP1

The KD values for the interaction between ACE2 and Fvc_1098 (23.4 nM) and Fvc_58 (4.48 μM) were significantly lower than for the interaction between ACE2 and Fvp (66.8 μM, Fig. 3A). This suggests that the binding affinity of crude fucoidan for ACE2 is much stronger than that of pure fucoidan and falls within similar nanomolar range as those previously observed for RBD-ACE2 interactions (5-75 nM) [10, 11, 52]. Of note, in our control samples binding affinity between Wuhan-Hu-1 RBD-Am-Biotin and ACE2 was in the lower nanomolar range (0.39-31.8 nM) (Supplementary Fig. 8A). Additionally, binding of Fvc to ACE2 was confirmed using the sandwich method for the highest tested Fvc concentration (Supplementary Fig. 8B). The association profile of the RBD ligand with the ACE2 analyte resulted in a 0.7 nm increase in the BLI response signal. Subsequently, exposure of the sensor to the Fvc analyte led to a further 0.5 nm increase in the response signal. These results support the hypothesis that the Fvc macromolecule can bind directly to ACE2 as well to the ACE2-RBD complex. Recently, Shi et al. [53] demonstrated that fucoidans from *A. nodosum* and *U. pinnatifida* significantly reduced ACE2 mRNA and protein expression *in vitro*, as well as in the lungs and GI tract of SARS-CoV-2 infected hamsters, indicating specific ACE2 inhibition, which reduced GI pathology, and enhanced the abundance of Bacteroidota and Patescibacteria. These findings demonstrate that fucoidans influence ACE2 activity through multiple mechanisms. The small intestine exhibits the second highest level of ACE2 expression, following the testis, with lower abundance in the stomach and large intestine [54, 55]. The presence of *Bacteroides dorei, B. thetaiotaomicron, B. massiliensis*, and *B. ovatus*, previously reported to reduce ACE2 expression in the murine gut (as cited in [56]), was inversely associated with SARS-CoV-2 levels in fecal samples from hospitalized COVID-19 patients [56]. Strategies aimed at modifying the intestinal microbiota by prophylactic and/or therapeutic fucoidan supplements could potentially reduce disease severity.

**Fig. 3.**
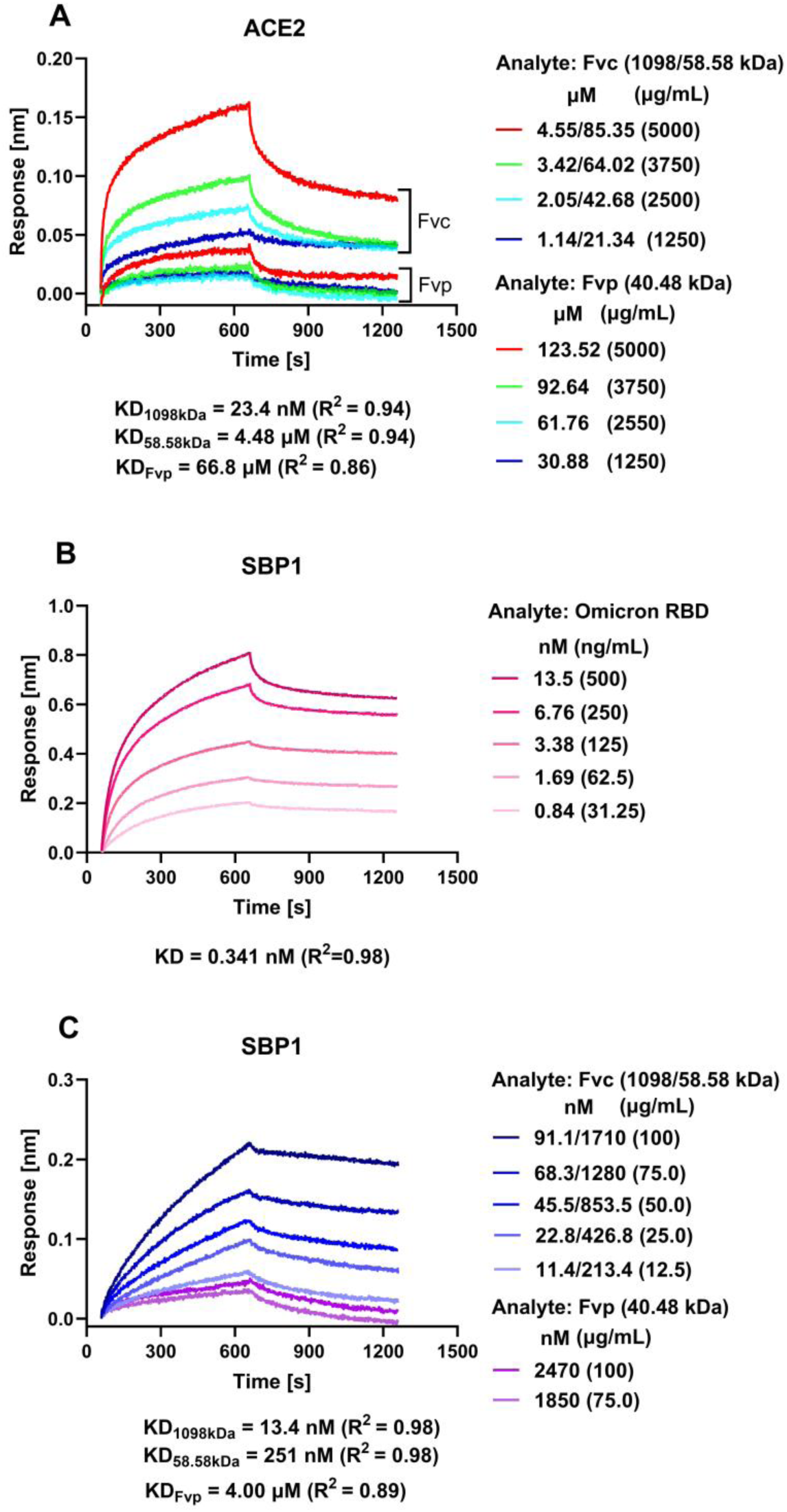
Sensorgrams of Fvc and Fvp binding to ACE2 (A), Omicron RBD to SBP1 (B), and Fvc and Fvp to SBP1(C).

We next evaluated the interaction between Fvc and SBP1, a 23-mer fragment derived from the α1-helix of the ACE2 peptidase domain. This peptide was identified through molecular dynamics simulations as a key element in ACE2 binding to the SARS-CoV-2 RBD [57]. Zhang et al. [57] also demonstrated that N-terminal biotinylated SBP1 binds with micromolar affinity (KD = 1.3 µM) to insect-derived SARS-CoV-2 RBD. In our study, SBP1 loaded on SAX biosensors showed much stronger binding affinity for Omicron RBD analyte (0.341 nM) (Fig. 3B). This discrepancy may, in part, be attributed to differences in the buffer composition during the BLI assays and to the type of biosensor used [58]. SBP1 exhibited stronger binding affinities for Fvc_1098 (KD = 13.4 nM) and Fvc_58 (KD = 251 nM) compared to Fvp (KD = 4.00 µM) (Fig. 3C).

### 3.5. SARS-CoV-2 pseudovirus entry inhibition assay

To determine the antiviral potency of Fvc and Fvp, a cell based functional assay was used. Inhibition of pseudovirus entry into HEK293T-ACE2-OE cells was observed in a dose-dependent manner across a range of concentrations (10 to 5000 μg/mL). Fvc demonstrated the most potent inhibitory effect (IC50 = 58 μg/mL), followed by Fvp (239 μg/mL) and alginate from *A. nodosum* (344 μg/mL) (Fig. 4A-B). In contrast, mannan, a non-sulfated polysaccharide, showed a less potent inhibitory effect, achieving approximately 60% inhibition at the highest concentration tested (5000 μg/mL). These results suggest that sulfated fucose-containing polysaccharides are more potent inhibitors. Additionally, fucoidans did not cause cytotoxicity in HEK293T cells, indicating the safety of these polysaccharides (Fig. 4C). Similarly, high molecular weight fucoidan (8.3 μg/mL) from *S. japonica* inhibited SARS-CoV-2 *in vitro* more effectively than low molecular weight fucoidan (16 μg/mL), heparin (36 μg/mL), and trisulfated-heparin (88 μg/mL) [59]. This enhanced activity was hypothesized to results from the greater multipoint binding capacity of the higher molecular weight fucoidan to the SARS-CoV-2 spike protein [59]. In contrast, unfractionated fucoidans from *F. vesiculosus* and *U. pinnatifida* were reported to be ineffective at inhibiting viral infection in Vero 76 cells [28]. However, Yang et al. [60] reported that both sulfated and desulfated fucoidans from *U. pinnatifida* and *F. vesiculosus* showed similar inhibitory effects on HCoV-OC43 and SARS-CoV-2 in HCT-8 and Vero E6 cells, respectively. The inhibitory concentrations observed in our study were comparable to those reported for fucoidans, from *A. nodosum* (48.66 μg/mL) and *U. pinnatifida* (69.52 μg/mL), which inhibited the entry of SARS-CoV-2 virus-like particles into Caco-2-N^int^ cells [53]. Consistent with our observations, He et al. [23] also demonstrated a high efficacy of naturally occurring, marine sulfated glycans in inhibiting the binding of both the wild type and XBB.1.5 RBDs to surface-immobilized heparin. Chemically desulfated glycans, demonstrated decreased binding activity, emphasizing the importance of sulfation for the inhibitory function of these glycans. In addition, Song et al. [18] reported that the IC50 values of the naturally derived unfractionated fucoidan from *S. japonica* (0.52 μg/mL) and rhamnan sulfate from *Monostroma nitidum* (0.93 μg/mL) were significantly lower than those of semi-synthetic pentosan polysulfate (1.6 μg/mL) and heparin (4.6 μg/mL). These findings suggest that the high potency observed for Fvc may be attributed to the preservation of its native structural characteristics during partial purification, including its high molecular weight and high fucose content. As Fvc sample contains traces of uronic acid [61–63], alginates with a molecular weight of 100 kDa from *F. vesiculosus* (AlgFv) and 300 kDa from *A. nodosum* (AlgAn) were used to study their binding to the Wuhan-Hu-1-RBD-Am-Biotin protein. The high KD value (7590 µM) for RBD-AlgFv interaction indicates weak affinity. These data suggest that the chemical moieties required for the interaction between fucoidan-RBD and alginate-RBD are distinct [64]. In contrast, the KD (2.61 µM) for RBD-AlgAn was comparable to that of RBD-Fvc_58 (4.27µM) (Supplementary Fig. 9). Other studies have reported that the crude alginate-containing polysaccharide and low molecular weight alginate derivatives inhibited spike protein binding to ACE2, with IC50 values ranging from 85.5 nM to 1.42 μM [60, 65]. The residues within the RBD crucial for interaction with high molecular weight alginates, which are negatively charged, remain unknown [64].

**Fig. 4.**
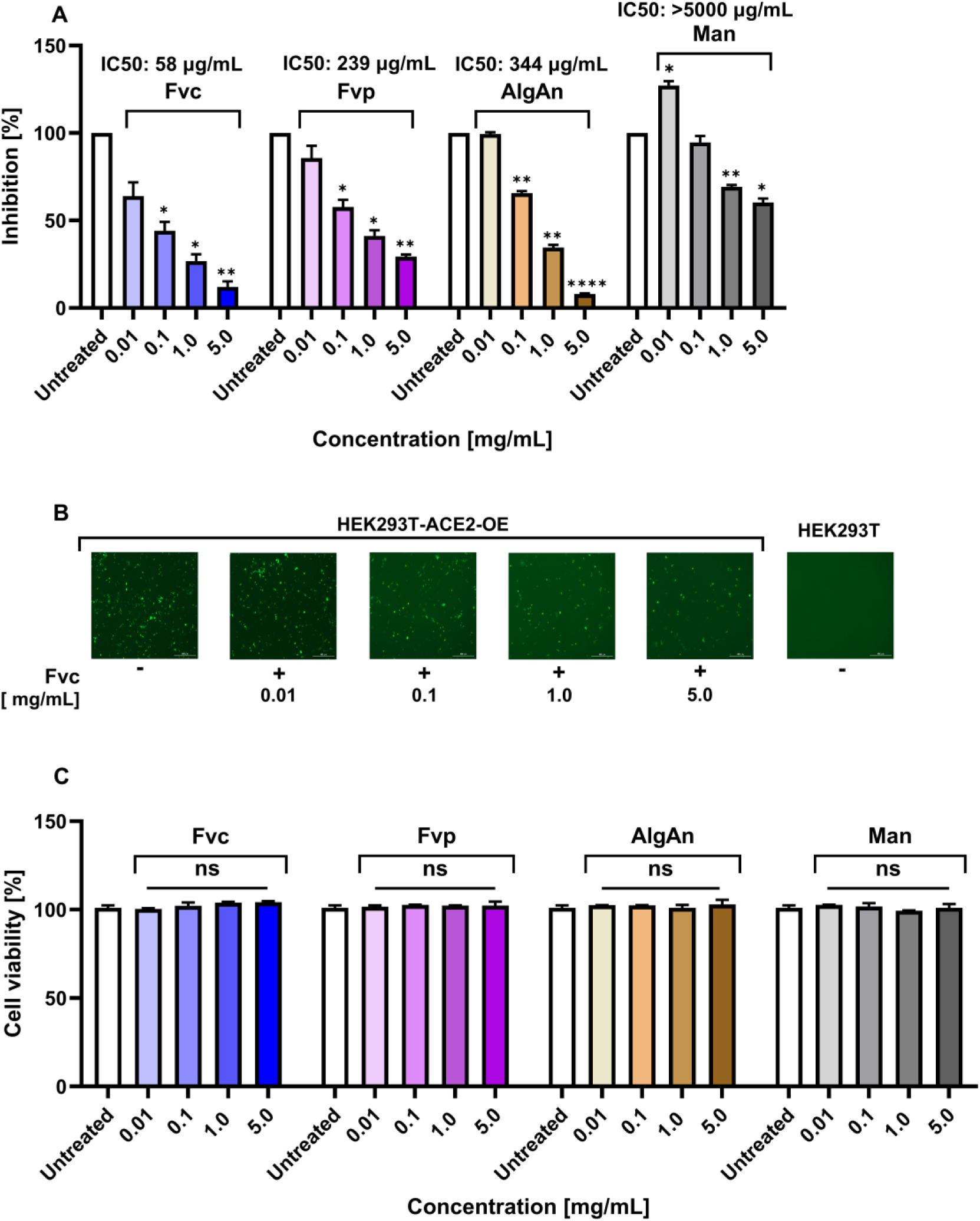
Effect of Fvc and Fvp on pseudovirus infection of HEK293T-ACE2-OE cells. Dose dependent entry inhibition by Fvc and Fvp (A), GFP positive cells in the representative images of pseudovirus entry (B), Cell viability in HEK293T-ACE2-OE cells by WST-1 assay (C). A two-way repeated-measures ANOVA followed by Bonferroni’s multiple comparisons test; ns, not significant (p ≥ 0.05); * p < 0.05; ** p < 0.01; **** p < 0.0001 denote differences vs. untreated controls. Additional pairwise differences between inhibitors and concentrations are summarized in Supplementary Table 3.

## 4. Conclusion

The results show that crude fucoidan Fvc has strong binding affinity for SARS-CoV-2 RBDs. Two molecular weight populations and high fucose content were observed in its sample. This suggests that the molecular weight and composition of fucoidan are crucial for its antiviral activity. The binding affinities for Fvc fall within the expected moderate (µM) to strong (nM) range for carbohydrate-protein interactions. Fvc generally exhibits better binding affinity than Fvp for both RBD and ACE2. KD values for both targets indicate a dual-target interaction. At nanomolar concentrations of Fvc and Fvp, binding was observed only to the Avi-tag-, but not amine-biotinylated RBD proteins. This confirms that single-side Avi-tag vs. random amine protein biotinylation enables better accessibility to its binding sites. This study provides strong evidence that crude, fucose rich and high molecular weight fucoidan from *F. vesiculosus* holds significant promise as an antiviral agent against SARS-CoV-2 as it outperformed pure fucoidan. Further research is needed to confirm whether the higher response signal for Fvc compared to Fvp, observed at micromolar concentrations but absent at nanomolar concentrations, is due to its high molecular weight fraction, aggregate formation, or the presence of unknown bioactive components that may be diminished at lower concentrations. Furthermore, fucoidan exhibited minimal cytotoxicity, suggesting that it could be a safe prophylactic and therapeutic option for SARS-CoV-2 and ACE2 dependent viral infections. High-dose oral administration of antiviral fucoidan as prophylactic or therapeutic agent could potentially reduce GI complications and fecal-oral route of viral transmission. However, additional studies, including pre-clinical mouse models are essential to fully assess its efficacy and ability to target emerging variants of the virus. Future research should also explore the potential synergistic effects of fucoidan in combination with other antiviral agents to improve treatment outcomes for patients.

## Acknowledgments

This work was financially supported in part by grants from the National Science and Technology Council, Taiwan (NSTC 111-2811-B-182A-018, 112-2811-B-182A-026, 113-2811-B-182A-026, 109-2314-B-182A-103-MY3, 112-2314-B-182A-070-MY3, and 114-2321-B-182-001) and from Chang Gung Memorial Hospital, Taoyuan, Taiwan (CMRPG3N1261-2, CORPG3K0311, CORPG3L0371, CORPG3M0411, and OMRPG3P0011). We thank all the staff members at core facilities for their technical support and expertise. The HPAEC-PAD and SEC-MALS analyses were supported by GRC Mass Core Facility of Genomics Research Center, and the Academia Sinica (AS) Glycoscience Core Facility (funded by the AS Core Facility and Innovative Instrument Project AS-CFII-114-A13), AS, Taipei, Taiwan. Biolayer Interferometry experiments were conducted with Octet RED96 instrument at the Core Instrument Center, High Throughput Analysis facility, Chang Gung University, Taoyuan, Taiwan. LC-MS/MS data were acquired at the Clinical Proteomics Core Laboratory of Chang Gung Memorial Hospital in Linkou, Taiwan. We thank Dr. Reiko Lee and Prof. Yuan-Chuan Lee (Department of Biology, Johns Hopkins University, Baltimore, Maryland, USA) for their valuable suggestions and comments on the ELISA, polysaccharide staining, and BLI assays.

## Declaration of competing interest

All authors declare no conflict of interest for this work and no financial and personal relationships with other people or organizations that could inappropriately influence or bias their work.

## Data availability

Data will be made available on request.

## Appendix A. Supplementary data

Supplementary data to this article can be found online

## Supplementary file 1

### 1. Materials and methods

#### 1.1. Enzyme-linked immunosorbent assays (ELISA)

Polysaccharides (100 µg/mL in 1 x PBS) such as fucoidans, alginates and dextrans (listed in the Supplementary Table 1) were coated overnight on Carbo-BIND ELISA microplates (Corning, NY, USA), then washed and blocked with PBSTB (1 x PBS + 0.05 % Tween 20 + 1 % bovine serum albumin (BSA)) [1]. Next, the plates were incubated with 2-fold serially diluted RBDs (10-0.078 µg/mL in PBSTB) for 60 min at room temperature at 100 rpm, followed by incubation with the anti-RBD antibody and HRP-conjugated secondary antibody (listed in the Supplementary Table 2) for 90 min. The reaction was developed using 3,3’,5,5’-tetramethylbenzidine substrate (TMB, NeA-Blue, Clinical Science Products, USA) for 30 min, and the color development was terminated by adding 1M H2SO4. Absorbance was measured at 450 nm. Experiments were performed in at least duplicates. Data are presented as mean ± SD (Supplementary Fig. 2A), with error bars representing SD (Supplementary Fig. 2B and C). To study polysaccharide inhibitory potency, first, the most potent fucoidan from the interaction assay, Fvc (100 µg/mL), was coated overnight at 4 °C on the Carbo-BIND ELISA microplates, and then blocked with PBSTB. Next, serially diluted polysaccharides (100-0.781 µg/mL) and RBD (1 µg/mL) were pre-incubated for 30 min and then added to fucoidan-coated wells [2]. The mixture was incubated for an additional 60 min and the RBDs detected as described above. Of note, neither heparin nor ACE2 could be used as an inhibition target, because heparin bound poorly to the plates and fucoidans showed binding to ACE2 (Fig. 3). IC50 values were estimated by using a free online program from AAT Bioquest (https://www.aatbio.com/tools/ic50-calculator).

#### 1.2. Fvc and Fvp proteomic analysis

Solutions of Fvc and Fvp, as well as the gel excised (at ∼ 100 kDa, PageRuler™ Prestained Protein Ladder, 10 to 180 kDa, # 26616, Thermo Fisher Scientific, USA) representative bands of Fvc and Fvp were analyzed by LC-MS/MS according to the modified method described by Feng et al. [3]. Tandem mass spectrometry analysis was performed on a Q-Exactive™ HF Mass Spectrometer (Thermo Fisher Scientific, San Jose, USA) coupled with a Thermo Scientific™ UltiMate™ 3000 RSLCnano high-pressure liquid chromatography (HPLC) system. The mixtures were directly loaded onto a 50-cm analytical column (EASY-Spray™ C18 Column) and separated by a 90-min linear gradient from 2 to 35 % buffer B (80 % acetonitrile in 0.1 % formic acid) at a flow rate of 200 nL/min.

The MS raw files were analyzed with MaxQuant version 1.5.3.30 using default settings. The minimal peptide length was set to 7. Trypsin was used as the digestion enzyme. Methylthio of cysteine was set as a fixed modification, while oxidation of methionine, pyroglutamic acid at the peptide N-terminus (Glutamine), and acetylation of the protein N-terminus were set as variable modifications. Up to two missed cleavages were allowed. The mass tolerance for the precursor was set to 20 ppm for the first search, 4.5 ppm for the main search, and 0.5 Da for the fragment ions. The files were searched against algal protein sequences of FASTA databases *Ectocarpus siliculosus*, *E. subulatus*, *Fucus*, *Porphyra umbilicalis*, *Saccharina japonica*, *Ulva mutabilis*, and *U. prolifera* [4–8]. Also, *Fucus vesiculosus* NCBI reference genome database was used. Protein identifications with at least one unique peptide were considered.

## Supplementary tables

**Table S1.**
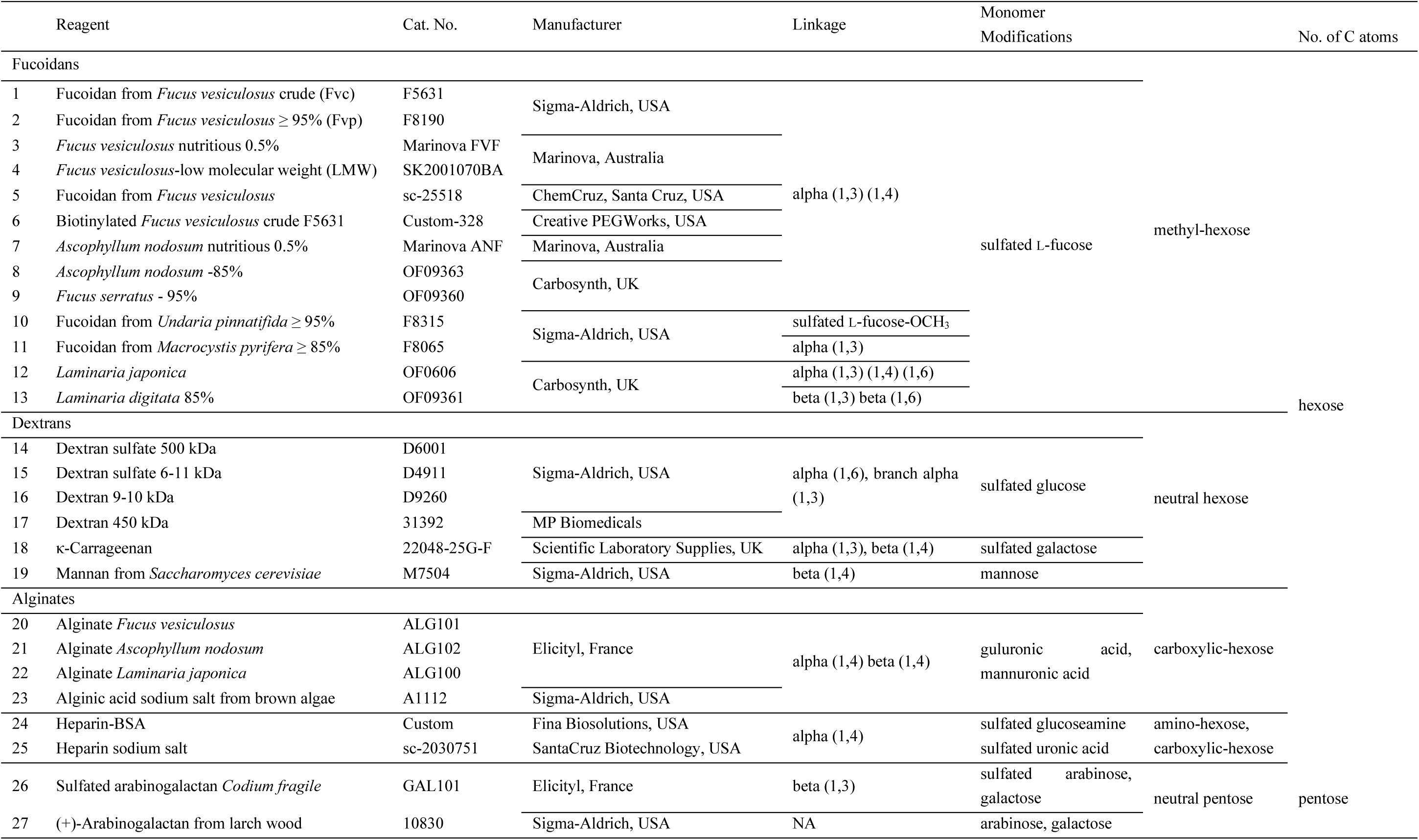
Polysaccharides used in this study.

**Table S2.**
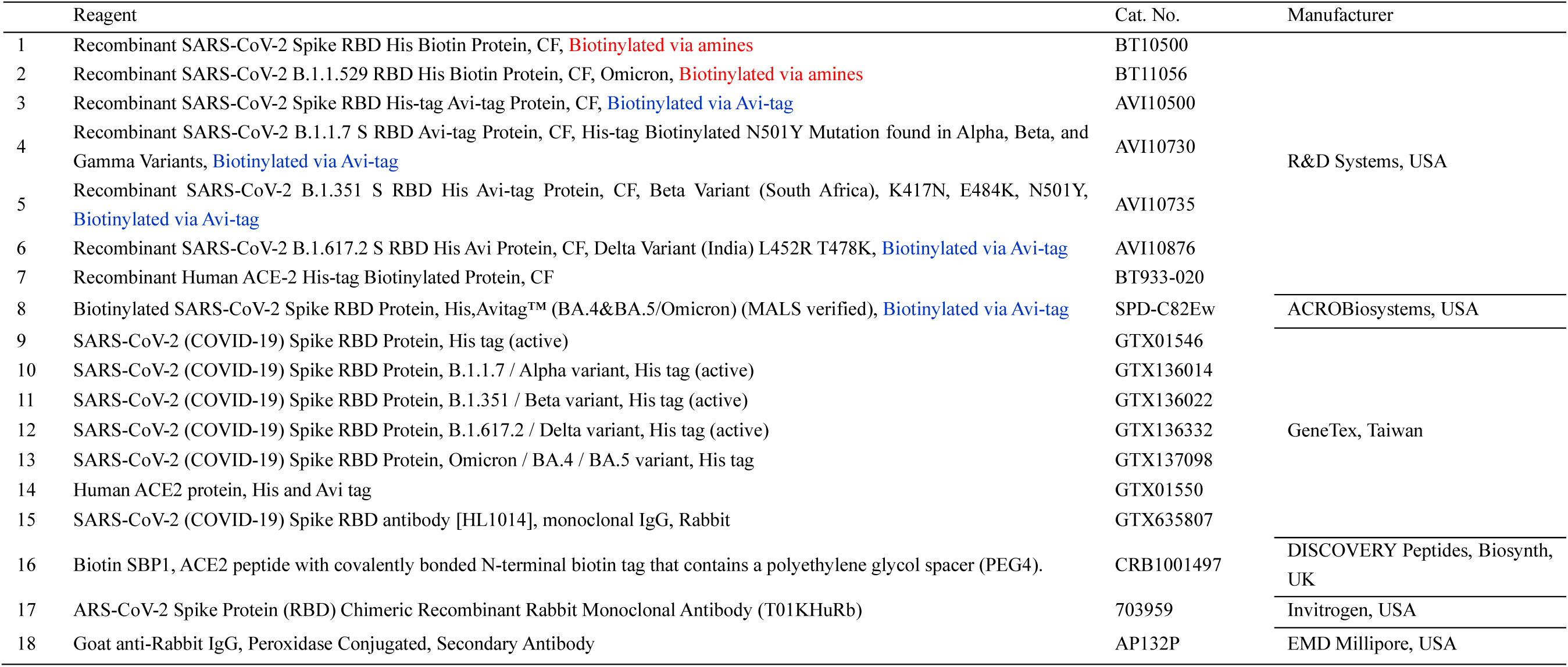
Proteins and peptides used in this study.

**Table S3.** Pairwise comparisons of % inhibition between inhibitors (A) and within each inhibitor across its concentrations (B) for the dose-dependent inhibition of pseudovirus entry into HEK293T-ACE2-OE cells. Bonferroni’s adjusted *p*-values were obtained from two-way repeated-measures ANOVA. Significance is indicated as: ns, not significant (p ≥ 0.05); * p < 0.05; ** p < 0.01; *** p < 0.001.

| Inhibitor/Concentration [mg/mL] | 0.01 | 0.1 | 1.0 | 5.0 |
| --- | --- | --- | --- | --- |
| Fvc vs. Fvp | 0.660 (ns) | 0.646 (ns) | 0.296 (ns) | 0.112 (ns) |
| Fvc vs. AlgAn | 0.269 (ns) | 0.256 (ns) | >0.999 (ns) | >0.999 (ns) |
| Fvc vs. Man | 0.060 (ns) | 0.010 (*) | 0.034 (*) | 0.002 (**) |
| Fvp vs. AlgAn | >0.999 (ns) | >0.999 (ns) | >0.999 (ns) | 0.009 (**) |
| Fvp vs. Man | 0.111 (ns) | 0.016 (*) | 0.050 (ns) | 0.008 (**) |
| AlgAn vs. Man | 0.020 (*) | 0.053 (ns) | <0.001 (***) | 0.009 (**) |

B. Within-inhibitor comparisons between concentrations.
| Concentration [mg/mL]/Inhibitor | Fvc | Fvp | AlgAn | Man |
| --- | --- | --- | --- | --- |
| 0.01 vs. 0.1 | 0.251 (ns) | 0.209 (ns) | 0.003 (**) | 0.062 (ns) |
| 0.01 vs. 1.0 | 0.130 (ns) | 0.130 (ns) | 0.008 (**) | 0.029 (*) |
| 0.01 vs. 5.0 | 0.091 (ns) | 0.125 (ns) | <0.001 (***) | 0.042 (*) |
| 0.1 vs. 1.0 | 0.040 (*) | 0.047 (*) | 0.025 (*) | 0.307 (ns) |
| 0.1 vs. 5.0 | 0.035 (*) | 0.114 (ns) | 0.003 (**) | 0.279 (ns) |
| 1.0 vs. 5.0 | 0.037 (*) | 0.297 (ns) | 0.037 (*) | 0.209 (ns) |

## Supplementary figures

**Fig. S1.**
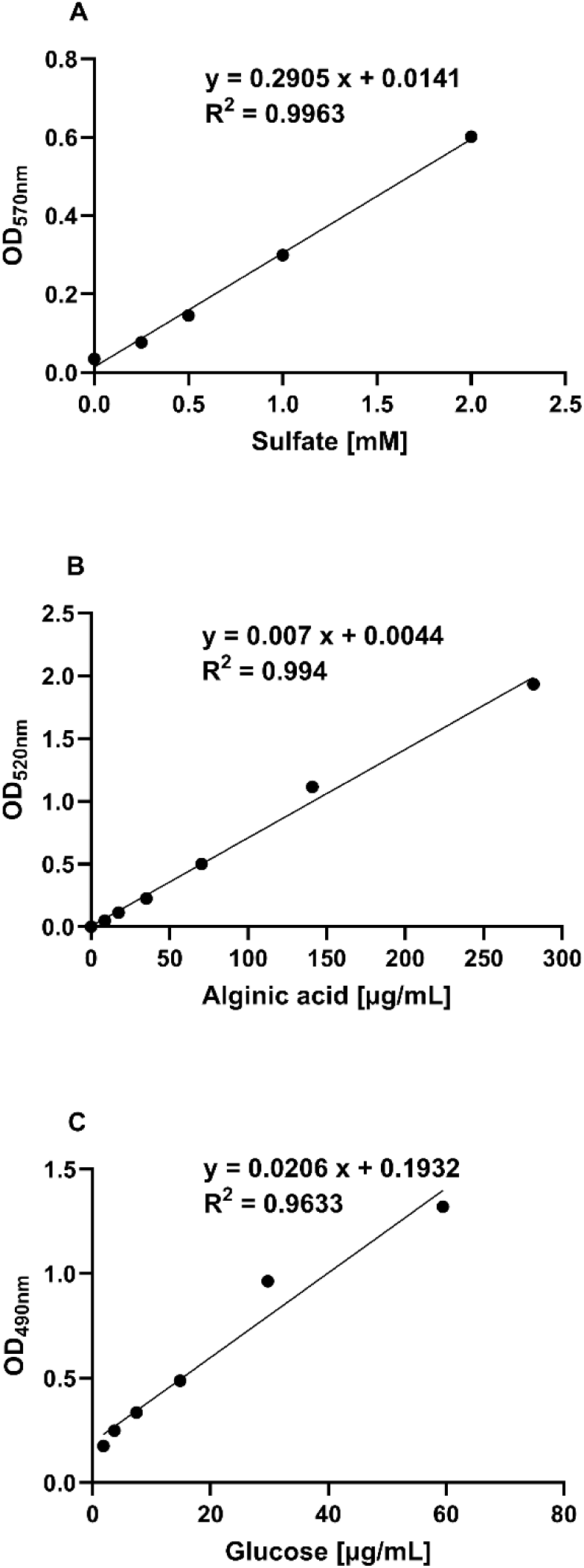
Standard curves for sulfate (A), uronic acid (B), and total carbohydrate (C) content.

**Fig. S2.**
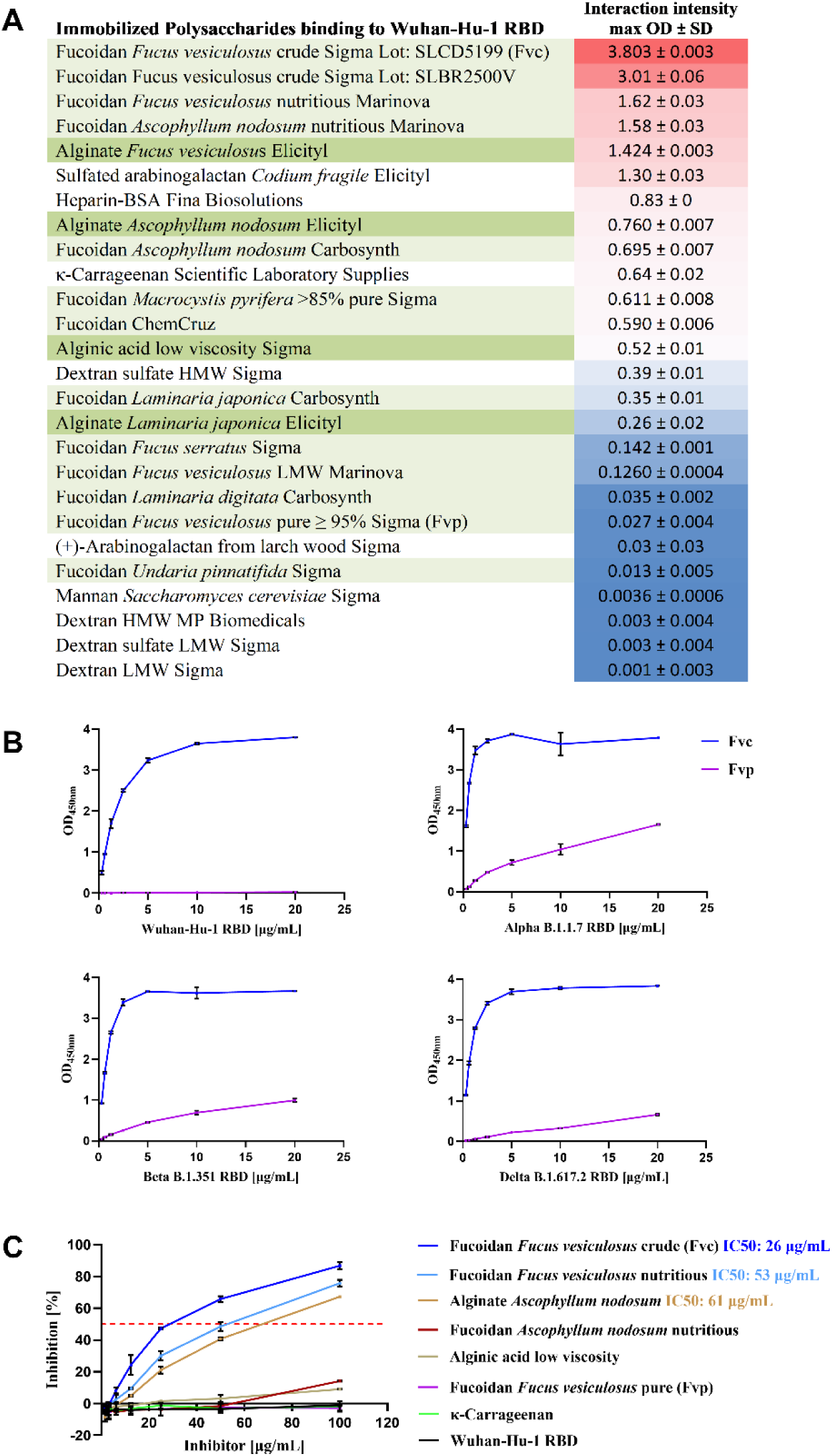
Biding (A, B) and inhibition (C) assays of RBDs to immobilized polysaccharides on Carbo-BIND-ELISA microplates. A color gradient heat map with high reactivity (red) to no reactivity (blue) of fucoidans (light green), alginates (dark green) and other polysaccharides.

**Fig. S3.**
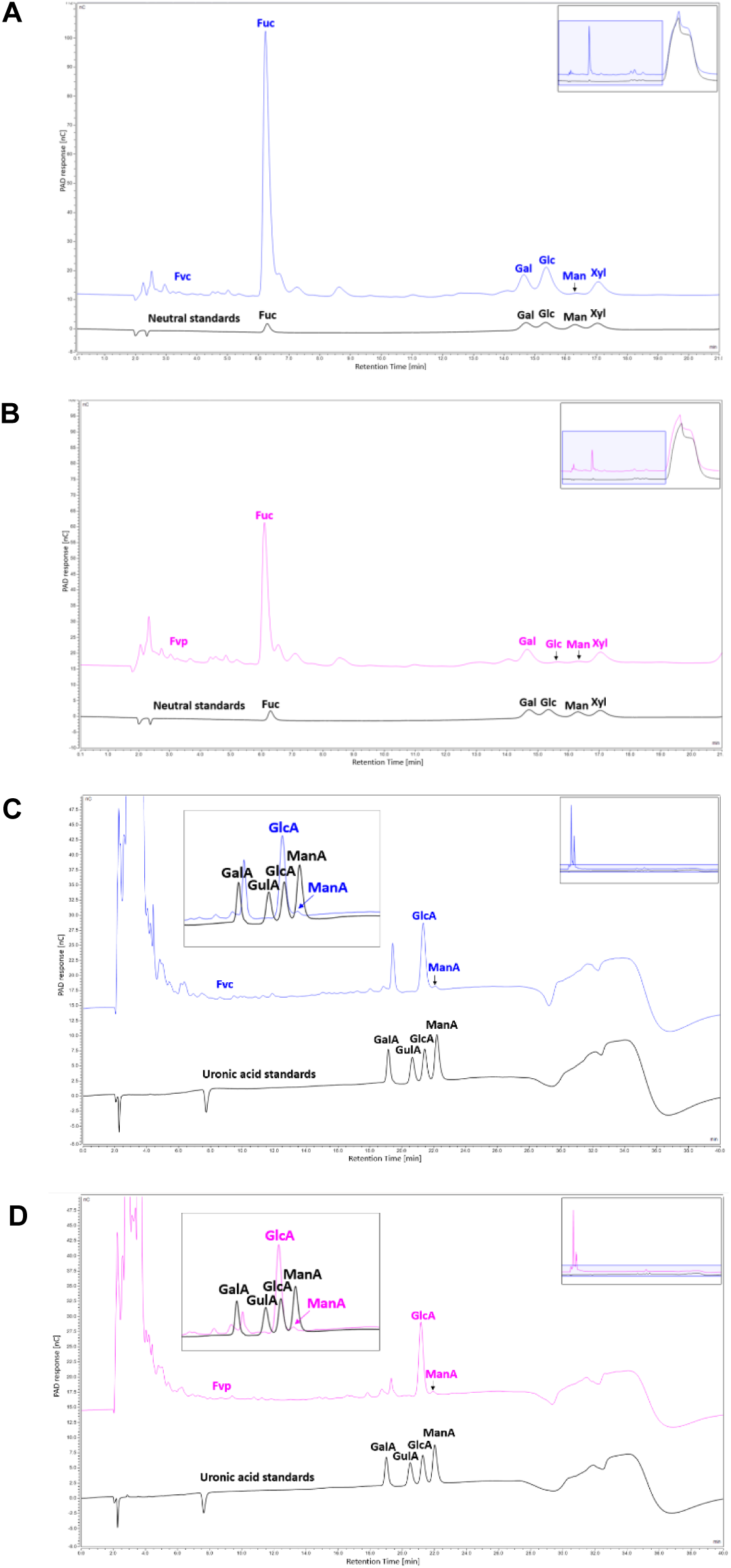
HPAEC-PAD chromatograms of neutral monosaccharides and uronic acid standards (black lines), Fvc (blue lines) (A, C), Fvp (pink lines) (B, D).

**Fig. S4.**
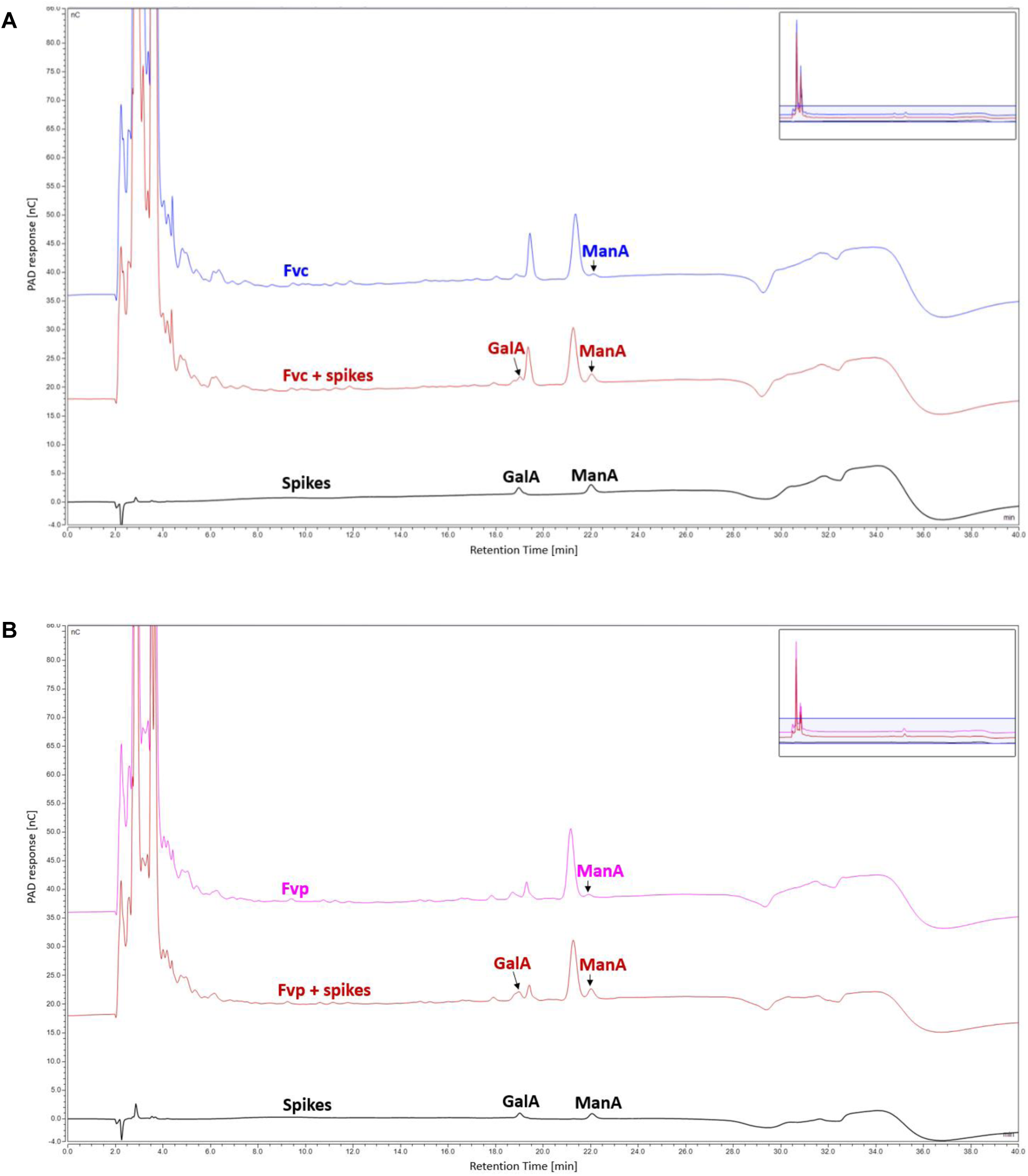
HPAEC-PAD chromatograms of uronic acid standards/spikes (black lines), Fvc (blue line) (A), Fvp (pink line) (B), Fvc and Fvp spiked with uronic acid standards (red lines).

**Fig. S5.**
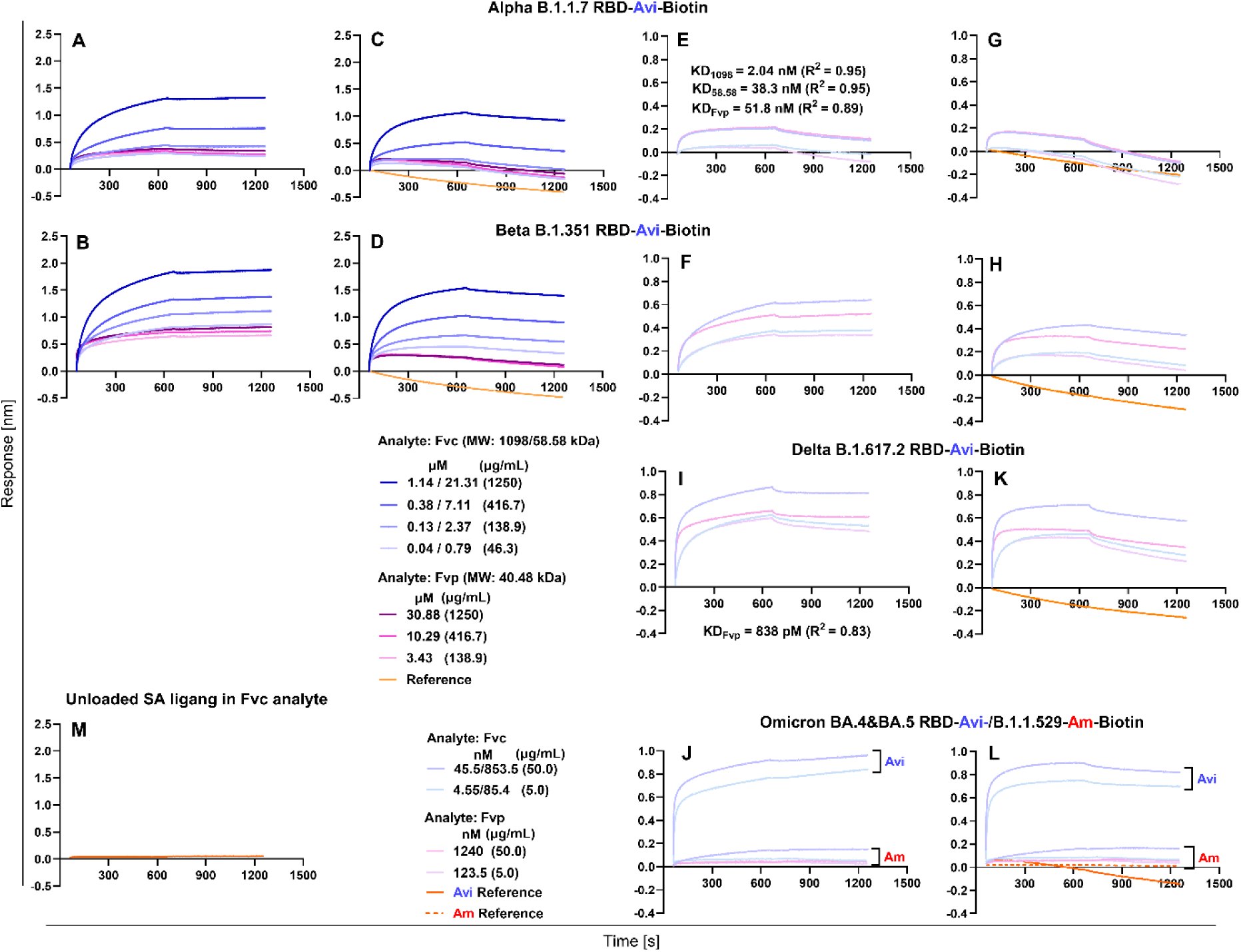
Sensorgrams of Fvc (blue lines) and Fvp (purple lines) binding to the Avitag- (Avi) and amine- (Am) biotinylated RBD ligands at micromolar concentration range of fucoidans (A) to (D), at nanomolar concertation range of fucoidans (E) to (L), unloaded SA biosensor in Fvc analyte (M).

**Fig. S6.**
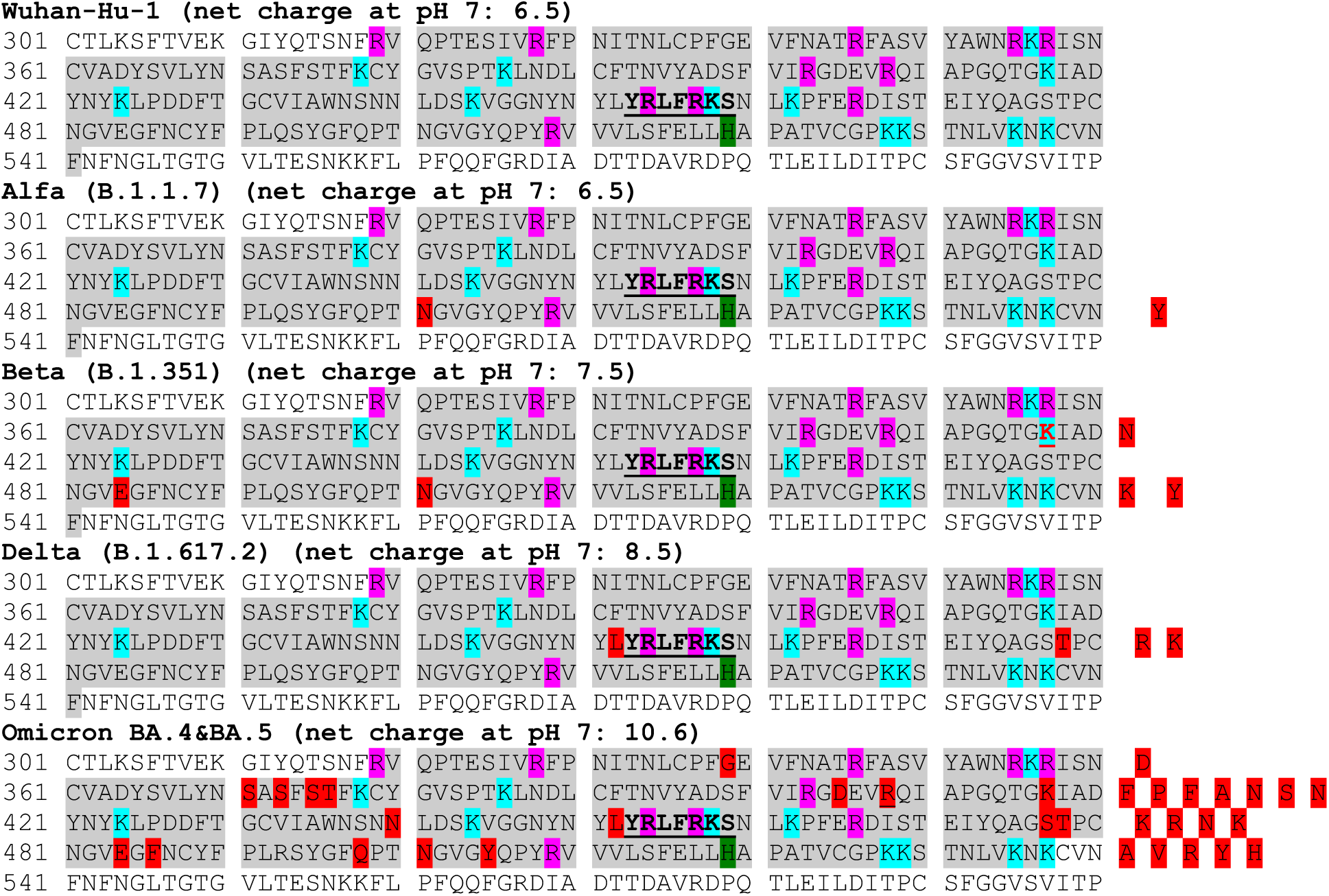
Recombinant protein sequences specific for RBD (R319-F541 aa for Wuhan-Hu-1, Alfa, Beta, and Delta, and R319-K537 aa for Omicron, highlighted in gray), accession no. YP_009724390.1/QHD43416.1 with mutations highlighted in red (Alfa: N501Y; Beta: K417N, E484K, and N501Y; Delta: L452R and T478K; Omicron: G339D, S371F, S373P, S375F, T376A, D405N, R408S, K417N, N440K, L452R, S477N, T478K, E484A, F486V, Q498R, N501Y, and Y505H), according to the information provided by the protein manufacturer (Supplementary Table 2). The GAG-binding motif (YRLFRKS) is underlined and bolded (453-459 aa). Positively charged amino acids histidine (H), lysine (K), arginine (R) are highlighted in green, blue, and pink, respectively. The net charge of each RBD sequence (highlighted in gray) at pH 7 was calculated with Innovagen Protein Calculator (http://pepcalc.com/protein-calculator.php).

**Fig. S7.**
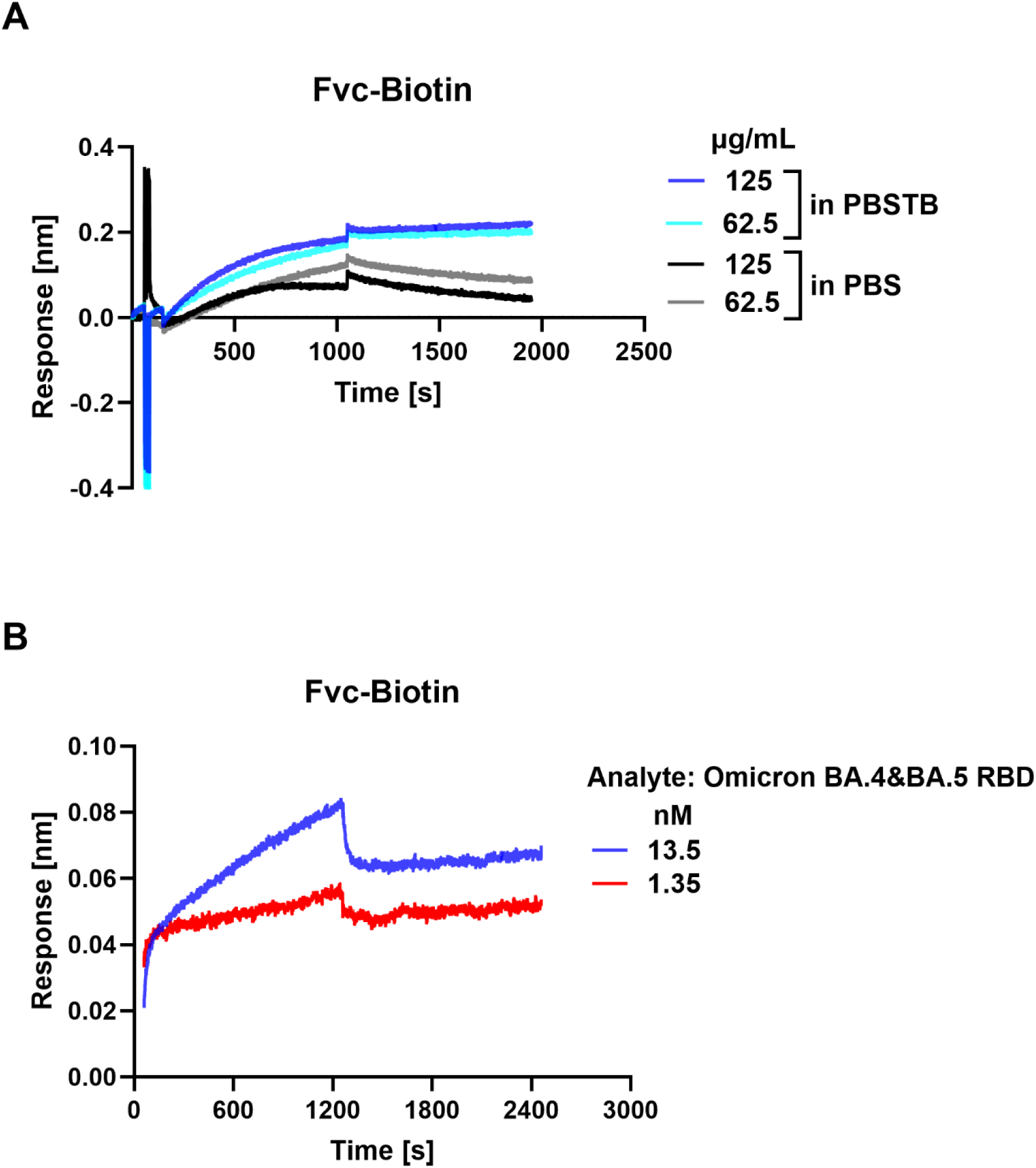
Loading of biotinylated Fvc (Fvc-Biotin) on SA biosensor (A), sensorgrams of BA.4&BA.5 RBD binding to Fvc-Biotin ligand (B).

**Fig. S8.**
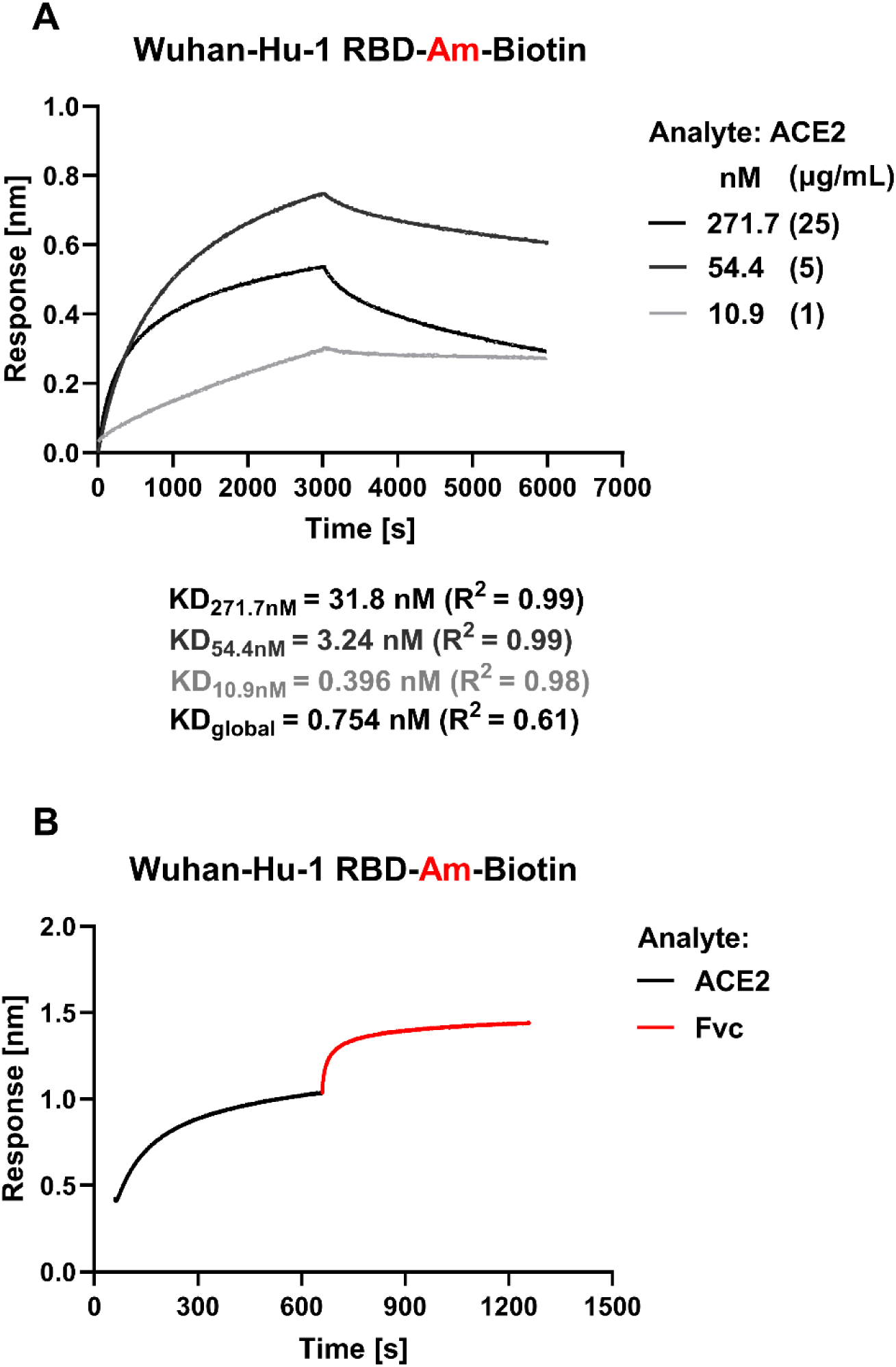
Sensorgrams of ACE2 binding to amine-biotinylated Wuhan-Hu-1 RBD (A), Fvc binding to ACE2 bound to amine biotinylated Wuhan-Hu-1 RBD ligand by the sandwich method (B).

**Fig. S9.**
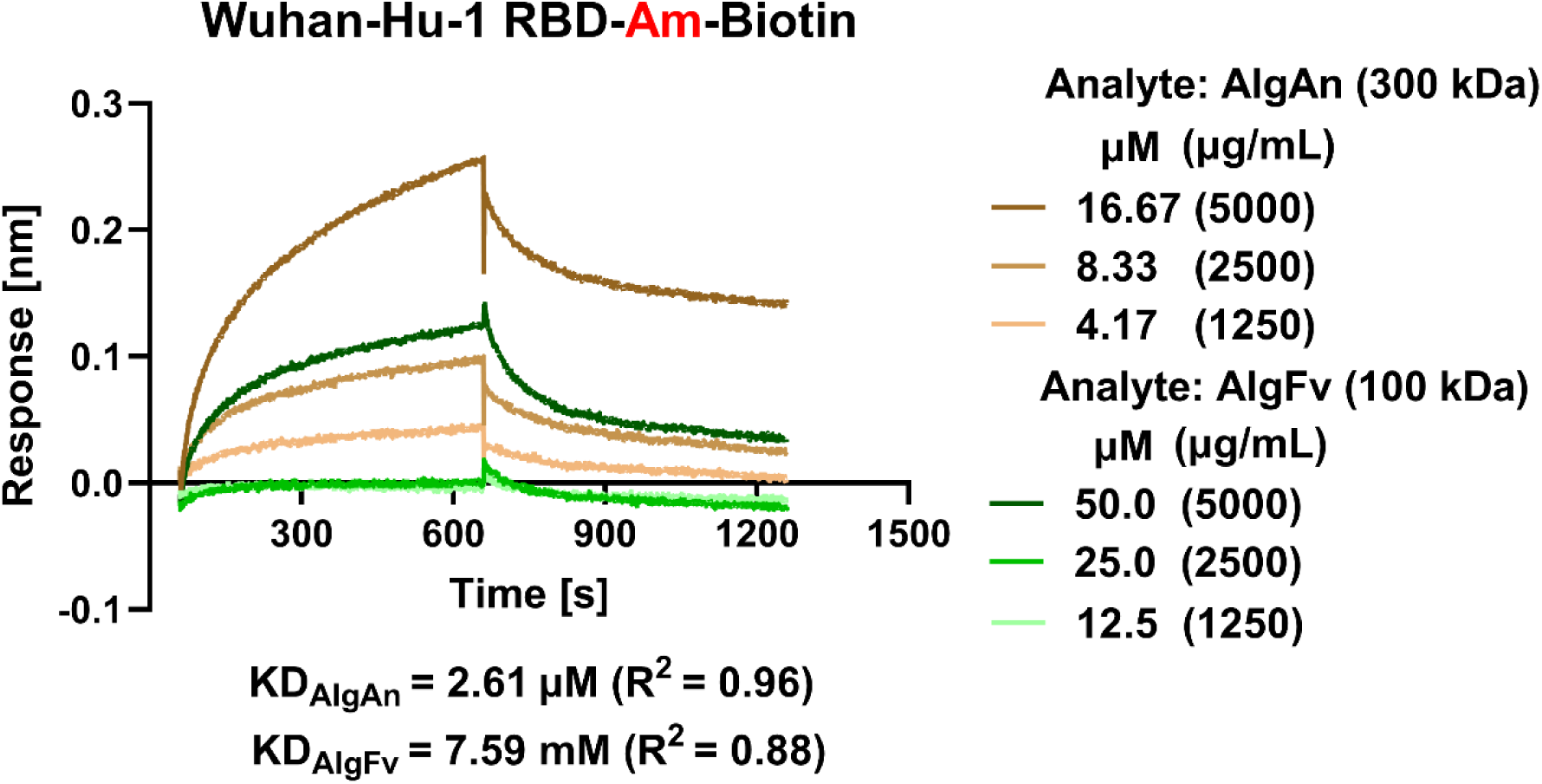
Sensorgrams of alginates from *F. vesiculosus* (AlgFv) and *A. nodosum* (AlgAn) binding to amine biotinylated Wuhan-Hu-1 RBD ligand.

## Supplementary file 2

Fvc_liquid

| **Fucus vesiculosus** |  |  |  |  |  |
| --- | --- | --- | --- | --- | --- |
| **Majority protein IDs** | **Protein names** | **Fasta headers** | **Unique peptides** | **Mol. weight [kDa]** | **iBAQ** |
| YP_005090070.1;CAX12450.1 | photosystem I iron-sulfur center (chloroplast) [Fucus vesiculosus] | YP_005090070.1 photosystem I iron-sulfur center (chloroplast) [Fucus vesiculosus];CAX12450.1 photosystem I iron-sulfur center (chloroplast) [Fucus vesiculosus] | 1 | 8.772 | 709910 |
| YP_005090031.1;CAX12411.1 | cytochrome c553 (chloroplast) [Fucus vesiculosus] | YP_005090031.1 cytochrome c553 (chloroplast) [Fucus vesiculosus];CAX12411.1 cytochrome c553 (chloroplast) [Fucus vesiculosus] | 1 | 12.094 | 703630 |
| YP_005090062.1;CAX12442.1;ABC25542.1;AA D43281.1;AAW70170.1 | ribulose-1,5-bisphosphate carboxylase/oxygenase large subunit (chloroplast) [Fucus vesiculosus] | YP_005090062.1 ribulose-1,5-bisphosphate carboxylase/oxygenase large subunit (chloroplast) [Fucus vesiculosus];CAX12442.1 ribulose-1,5-bisphosphate carboxylase/oxygenase large subunit (chloroplast) [Fucus vesiculosus];ABC25542.1 ribulose-1,5-bisphosphate c | 4 | 53.934 | 503700 |
| YP_005090091.1;CAX12474.1 | cytochrome b559 alpha chain (chloroplast) [Fucus vesiculosus] | YP_005090091.1 cytochrome b559 alpha chain (chloroplast) [Fucus vesiculosus];CAX12474.1 cytochrome b559 alpha chain (chloroplast) [Fucus vesiculosus] | 2 | 9.5896 | 464410 |
| sp\|Q39757.1\|1433_FUCVE;CAA67389.1 | 14-3-3-like protein [Fucus vesiculosus] | sp\|Q39757.1\|1433_FUCVE RecName: Full=14-3-3-like protein;CAA67389.1 14-3-3 [Fucus vesiculosus] | 2 | 28.169 | 413060 |
| YP_005090114.1;CAX12500.1 | photosystem II reaction center protein D2 (chloroplast) [Fucus vesiculosus] | YP_005090114.1 photosystem II reaction center protein D2 (chloroplast) [Fucus vesiculosus];CAX12500.1 photosystem II reaction center protein D2 (chloroplast) [Fucus vesiculosus] | 3 | 38.991 | 162600 |
| YP_005090030.1;CAX12410.1 | PsbV (chloroplast) [Fucus vesiculosus] | YP_005090030.1 PsbV (chloroplast) [Fucus vesiculosus];CAX12410.1 PsbV (chloroplast) [Fucus vesiculosus] | 1 | 17.712 | 116640 |
| YP_005090063.1;CAX12443.1 | ribulose-1,5-bisphosphate carboxylase/oxygenase small subunit (chloroplast) [Fucus vesiculosus] | YP_005090063.1 ribulose-1,5-bisphosphate carboxylase/oxygenase small subunit (chloroplast) [Fucus vesiculosus];CAX12443.1 ribulose-1,5-bisphosphate carboxylase/oxygenase small subunit (chloroplast) [Fucus vesiculosus] | 1 | 16.006 | 105600 |
| sp\|Q39758.1\|ACT_FUCVE;CAA67388.1 | Actin; beta-actin [Fucus vesiculosus] | sp\|Q39758.1\|ACT_FUCVE RecName: Full=Actin;CAA67388.1 beta-actin [Fucus vesiculosus] | 2 | 41.889 | 80673 |
| YP_005090055.1;CAX12435.1 | ATP synthase CF1 alpha chain (chloroplast) [Fucus vesiculosus] | YP_005090055.1 ATP synthase CF1 alpha chain (chloroplast) [Fucus vesiculosus];CAX12435.1 ATP synthase CF1 alpha chain (chloroplast) [Fucus vesiculosus] | 1 | 54.095 | 59410 |
| AGK85255.1 | vacuolar ATPase subunit A [Fucus vesiculosus] | AGK85255.1 vacuolar ATPase subunit A [Fucus vesiculosus] | 1 | 68.399 | 19627 |
| **Ectocarpus siliculosus, Ectocarpus subulatus, Fucus, Porphyra umbilicalis, Saccharina japonica, Ulva mutabilis, Ulva prolifera** |  |  |  |  |  |
| CBN74826.1 | conserved unknown protein | CBN74826.1 conserved unknown protein [Ectocarpus siliculosus] | 1 | 23.531 | 1761900 |
| CBN77006.1;CBJ29487.1;CBJ32222.1;CBJ33691.1;CBJ33692.1;CBJ28877.1;CBJ29489.1;CBJ33060.1;CBN80506.1;CBN80505.1;CBJ33313.1;CBJ32234.1;CBJ32221.1;CBJ32149.1;ACE80197.1;ABB54384.1;CBJ33312.1;CBJ32225.1;CBJ34095.1;CBN80504.1;CBN80503.1;CBJ26059.1;CBJ32223.1;CBN80510.1 | Light harvesting complex protein | CBN77006.1 Light harvesting complex protein [Ectocarpus siliculosus];CBJ29487.1 Light harvesting complex protein [Ectocarpus siliculosus];CBJ32222.1 Light harvesting complex protein [Ectocarpus siliculosus];CBJ33691.1 Light harvesting complex protein [Ecto | 1 | 11.537 | 1295500 |
| CAA67389.1;sp\|Q39757.1\|1433_FUCVE;ACF06189.1 | 14-3-3 | CAA67389.1 14-3-3 [Fucus vesiculosus];sp\|Q39757.1\|1433_FUCVE RecName: Full=14-3-3-like protein;ACF06189.1 14-3-3-like protein, partial [Fucus serratus] | 2 | 28.169 | 413060 |
| OSX80310.1;OSX80297.1;CBN77459.1;CBN77456.1;CBN77451.1;CBJ48890.1;CBJ48889.1;CBJ32081.1;CBJ32073.1;CBJ33198.1 | hypothetical protein BU14_0055s0032 | OSX80310.1 hypothetical protein BU14_0055s0032 [Porphyra umbilicalis];OSX80297.1 hypothetical protein BU14_0055s0020 [Porphyra umbilicalis];CBN77459.1 histone H4 [Ectocarpus siliculosus];CBN77456.1 histone H4 [Ectocarpus siliculosus];CBN77451.1 histone H4 | 1 | 11.483 | 412250 |
| CBN77458.1;CBN77452.1;CBN77409.1;CBN74672.1;CBJ48886.1;CBJ33202.1;CBJ32076.1 | conserved unknown protein | CBN77458.1 conserved unknown protein [Ectocarpus siliculosus];CBN77452.1 conserved unknown protein [Ectocarpus siliculosus];CBN77409.1 conserved unknown protein [Ectocarpus siliculosus];CBN74672.1 conserved unknown protein [Ectocarpus siliculosus];CBJ48886 | 2 | 12.798 | 365430 |
| YP_005090091.1;CAX12474.1;AVZ00611.1 | cytochrome b559 alpha chain (chloroplast) | YP_005090091.1 cytochrome b559 alpha chain (chloroplast) [Fucus vesiculosus];CAX12474.1 cytochrome b559 alpha chain (chloroplast) [Fucus vesiculosus];AVZ00611.1 cytochrome b559 alpha chain (plastid) [Fucus spiralis] | 1 | 9.5896 | 301610 |
| CBJ26867.1 | COX15 homolog, cytochrome c oxidase assembly protein | CBJ26867.1 COX15 homolog, cytochrome c oxidase assembly protein [Ectocarpus siliculosus] | 1 | 40.016 | 295090 |
| CBN77947.1 | profilin | CBN77947.1 profilin [Ectocarpus siliculosus] | 1 | 16.453 | 281780 |
| CBJ26193.1 | LRR-GTPase of the ROCO family | CBJ26193.1 LRR-GTPase of the ROCO family [Ectocarpus siliculosus] | 1 | 136.61 | 280160 |
| CBJ27783.1 | imm downregulated 23 | CBJ27783.1 imm downregulated 23 [Ectocarpus siliculosus] | 1 | 19.67 | 210070 |
| WYC12533.1;WYC12531.1;WYC12530.1;CBJ30601.1;ACJ05913.1;ABB80121.1;ABB02445.1;sp\|Q39758.1\|ACT_FUCVE;CAA67388.1;ARO49595.1;ARO49594.1 | actin | WYC12533.1 actin [Saccharina japonica];WYC12531.1 actin [Saccharina japonica];WYC12530.1 actin [Saccharina japonica];CBJ30601.1 actin [Ectocarpus siliculosus];ACJ05913.1 beta-actin [Saccharina japonica];ABB80121.1 actin [Saccharina japonica];ABB02445.1 act | 2 | 41.79 | 187880 |
| ABC25542.1;YP_005090062.1;CAX12442.1;AVZ00582.1;WGS92535.1;WKW95982.1;CBJ55517.1;AAD43281.1;AAW70170.1;WGS91979.1;YP_006639053.1;YP_003289289.1;WKW96095.1;AAP78676.1;AAD41743.1;WGS92257.1;WDS74816.1;ADM16563.1;WAX38128.1;QIE12374.1;CBH31935.1;CAV31320.1;CAT18876.1;AGM75439.1;AFC40107.1;WKW95843.1 | ribulose-1,5-bisphosphate carboxylase/oxygenase large subunit (chloroplast) | ABC25542.1 ribulose-1,5-bisphosphate carboxylase/oxygenase large subunit (chloroplast) [Fucus vesiculosus];YP_005090062.1 ribulose-1,5-bisphosphate carboxylase/oxygenase large subunit (chloroplast) [Fucus vesiculosus];CAX12442.1 ribulose-1,5-bisphosphate c | 3 | 53.934 | 178270 |
| CBJ28399.1 | imm downregulated 8 | CBJ28399.1 imm downregulated 8 [Ectocarpus siliculosus] | 1 | 26.491 | 169110 |
| YP_006639054.1;YP_005090063.1;YP_003289290.1;WKW96150.1;WKW95983.1;WKW95844.1;WGS92536.1;WGS92258.1;WGS91980.1;WDS74817.1;WAX38129.1;sp\|P24395.1\|RBS_ECTSI;QIE12375.1;CAX12443.1;CAV31321.1;C | ribulose-1,5-bisphosphate carboxylase/oxygenase small subunit (chloroplast) | YP_006639054.1 ribulose-1,5-bisphosphate carboxylase/oxygenase small subunit (chloroplast) [Saccharina japonica];YP_005090063.1 ribulose-1,5-bisphosphate carboxylase/oxygenase small subunit (chloroplast) [Fucus vesiculosus];YP_003289290.1 Ribulose bisphosph | 1 | 15.836 | 118790 |
| AT18877.1;CAA36744.1;AVZ00583.1;AFC40108.1;ADM16564.1 |  |  |  |  |  |
| CBN78618.1 | conserved unknown protein | CBN78618.1 conserved unknown protein [Ectocarpus siliculosus] | 1 | 71.094 | 114750 |
| OSX75295.1 | hypothetical protein BU14_0242s0013 | OSX75295.1 hypothetical protein BU14_0242s0013 [Porphyra umbilicalis] | 1 | 42.002 | 111730 |
| OSX75725.1;CBJ32894.1;CBJ32893.1;OSX75723.1;OSX74984.1 | hypothetical protein BU14_0224s0023 | OSX75725.1 hypothetical protein BU14_0224s0023 [Porphyra umbilicalis];CBJ32894.1 EEF1A1, eukaryotic translation elongation factor 1 alpha [Ectocarpus siliculosus];CBJ32893.1 EEF1A2, eukaryotic translation elongation factor 1 alpha [Ectocarpus siliculosus]; | 1 | 41.84 | 109120 |
| CBN79094.1 | Trans-2-enoyl-CoA reductase, mitochondrial precursor | CBN79094.1 Trans-2-enoyl-CoA reductase, mitochondrial precursor [Ectocarpus siliculosus] | 1 | 38.246 | 87510 |
| CBN74529.1;CBN74283.1 | hypothetical protein Esi_0028_0170 | CBN74529.1 hypothetical protein Esi_0028_0170 [Ectocarpus siliculosus];CBN74283.1 conserved unknown protein [Ectocarpus siliculosus] | 1 | 72.438 | 68370 |
| CBJ32298.1 | F-type H-ATPase beta subunit | CBJ32298.1 F-type H-ATPase beta subunit [Ectocarpus siliculosus] | 1 | 53.838 | 59610 |
| OSX81338.1 | hypothetical protein BU14_0022s0058 | OSX81338.1 hypothetical protein BU14_0022s0058 [Porphyra umbilicalis] | 1 | 109.62 | 42299 |
| CBJ29307.1;OSX76008.1;CBJ48460.1;AEN71225.1;ACJ05912.1;CBJ29295.1;CBJ32839.1;OSX71346.1;CBN79394.1 | Heat shock protein 70 | CBJ29307.1 Heat shock protein 70, likely pseudogene [Ectocarpus siliculosus];OSX76008.1 hypothetical protein BU14_0212s0028 [Porphyra umbilicalis];CBJ48460.1 Molecular chaperones GRP78/BiP/KAR2, HSP70 superfamily [Ectocarpus siliculosus];AEN71225.1 heat sh | 1 | 66.577 | 41092 |
| CBN79008.1;AGK85255.1 | conserved unknown protein | CBN79008.1 conserved unknown protein [Ectocarpus siliculosus];AGK85255.1 vacuolar ATPase subunit A [Fucus vesiculosus] | 1 | 67.996 | 21310 |
| CBN79119.1 | conserved unknown protein | CBN79119.1 conserved unknown protein [Ectocarpus siliculosus] | 1 | 286.7 | 13680 |
| OSX79348.1 | LOW QUALITY PROTEIN: hypothetical protein BU14_0081s0027 | OSX79348.1 LOW QUALITY PROTEIN: hypothetical protein BU14_0081s0027 [Porphyra umbilicalis] | 1 | 300.93 | 13586 |
| YP_009413368.1;YP_006639138.1;YP_005090114.1;YP_003289147.1;WKW96107.1;WKW96067.1;WKW95928.1;WGS92620.1;WGS92342.1;WGS92064.1;WDS74901.1;WAX38213.1;QIE12379.1;CAX12500.1;CAV31178.1;CAT18680.1;AVZ00634.1;ASN78829.1;AFC40192.1;AFC40025.1 | photosystem II protein D2 (chloroplast) | YP_009413368.1 photosystem II protein D2 (chloroplast) [Porphyra umbilicalis];YP_006639138.1 photosystem II D2 protein (chloroplast) [Saccharina japonica];YP_005090114.1 photosystem II reaction center protein D2 (chloroplast) [Fucus vesiculosus];YP_0032891 | 1 | 39.086 | 0 |

Fvp\_liquid
| Fucus vesiculosus |  |  |  |  |  |
| --- | --- | --- | --- | --- | --- |
| Majority protein IDs | Protein names | Fasta headers | Unique peptides | Mol. weight [kDa] | iBAQ |
| YP_005090083.1;CAX12466.1 | photochlorophyllide reductase subunit L (chloroplast) [Fucus vesiculosus] | YP_005090083.1 photochlorophyllide reductase subunit L (chloroplast) [Fucus vesiculosus];CAX12466.1 photochlorophyllide reductase subunit L (chloroplast) [Fucus vesiculosus] | 1 | 32.228 | 1078100 |
| YP_005090131.1;CAX12517.1 | ribosomal protein L31 (chloroplast) [Fucus vesiculosus] | YP_005090131.1 50S ribosomal protein L31 (chloroplast) [Fucus vesiculosus];CAX12517.1 50S ribosomal protein L31 (chloroplast) [Fucus vesiculosus] | 1 | 8.2223 | 354440 |
| YP_005090065.1;CAX12445.1 | photosystem I P700 apoprotein A2 (chloroplast) [Fucus vesiculosus] | YP_005090065.1 photosystem I P700 apoprotein A2 (chloroplast) [Fucus vesiculosus];CAX12445.1 photosystem I P700 apoprotein A2 (chloroplast) [Fucus vesiculosus] | 1 | 82.494 | 77347 |
| sp\|Q39758.1\|ACT_FUCVE;CAA67388.1 | Actin; beta-actin [Fucus vesiculosus] | sp\|Q39758.1\|ACT_FUCVE RecName: Full=Actin;CAA67388.1 beta-actin [Fucus vesiculosus] | 1 | 41.889 | 49801 |
| YP_448626.1;AAR29320.1 | cytochrome c oxidase subunit 2 (mitochondrion) [Fucus vesiculosus] | YP_448626.1 cytochrome c oxidase subunit 2 (mitochondrion) [Fucus vesiculosus];AAR29320.1 cytochrome oxidase subunit II (mitochondrion) [Fucus vesiculosus] | 1 | 118.69 | 41675 |
| **Ectocarpus siliculosus, Ectocarpus subulatus, Fucus, Porphyra umbilicalis, Saccharina japonica, Ulva mutabilis, Ulva prolifera** |  |  |  |  |  |
| OSX78263.1;OSX69064.1;OSX77439.1;CBN79251.1;CBJ32017.1;OSX71664.1 | hypothetical protein BU14_0113s0008 | OSX78263.1 hypothetical protein BU14_0113s0008 [Porphyra umbilicalis];OSX69064.1 hypothetical protein BU14_1909s0002, partial [Porphyra umbilicalis];OSX77439.1 hypothetical protein BU14_0148s0015 [Porphyra umbilicalis];CBN79251.1 conserved unknown protein | 1 | 14.602 | 194860 |
| WYC12533.1;WYC12531.1;WYC12530.1;sp\|Q39758.1\|ACT_FUCVE;CBJ30601.1;CAA67388.1;ACJ05913.1;ABB80121.1;ABB02445.1 | actin | WYC12533.1 actin [Saccharina japonica];WYC12531.1 actin [Saccharina japonica];WYC12530.1 actin [Saccharina japonica];sp\|Q39758.1\|ACT_FUCVE RecName: Full=Actin;CBJ30601.1 actin [Ectocarpus siliculosus];CAA67388.1 beta-actin [Fucus vesiculosus];ACJ05913.1 be | 1 | 41.79 | 47429 |

Fvc\_gel
| **Fucus vesiculosus** |  |  |  |  |  |
| --- | --- | --- | --- | --- | --- |
| Majority protein IDs | Protein names | Fasta headers | Unique peptides | Mol. weight [kDa] | iBAQ |
| sp\|Q39758.1\|ACT_FUCVE;CAA67388.1 | Actin;CAA67388.1 beta-actin [Fucus vesiculosus] | sp\|Q39758.1\|ACT_FUCVE RecName: Full=Actin;CAA67388.1 beta-actin [Fucus vesiculosus] | 4 | 41.889 | 3548000 |
| sp\|Q39757.1\|1433_FUCVE;CAA67389.1 | 14-3-3-like protein [Fucus vesiculosus] | sp\|Q39757.1\|1433_FUCVE RecName: Full=14-3-3-like protein;CAA67389.1 14-3-3 [Fucus vesiculosus] | 2 | 28.169 | 2136700 |
| YP_448618.1;AAR29312.1 | ribosomal protein S11 (mitochondrion) [Fucus vesiculosus] | YP_448618.1 ribosomal protein S11 (mitochondrion) [Fucus vesiculosus];AAR29312.1 ribosomal protein S11 (mitochondrion) [Fucus vesiculosus] | 1 | 20.553 | 2026700 |
| YP_005090043.1;CAX12423.1 | photosystem I ferredoxin-binding protein (chloroplast) [Fucus vesiculosus] | YP_005090043.1 photosystem I ferredoxin-binding protein (chloroplast) [Fucus vesiculosus];CAX12423.1 photosystem I ferredoxin-binding protein (chloroplast) [Fucus vesiculosus] | 2 | 15.035 | 1874300 |
| YP_005090057.1;CAX12437.1 | putative peroxiredoxin ycf42 (chloroplast) [Fucus vesiculosus] | YP_005090057.1 putative peroxiredoxin ycf42 (chloroplast) [Fucus vesiculosus];CAX12437.1 putative peroxiredoxin ycf42 (chloroplast) [Fucus vesiculosus] | 1 | 21.202 | 1696600 |
| CAC36148.1 | ALA dehydratase [Fucus vesiculosus] | CAC36148.1 ALA dehydratase [Fucus vesiculosus] | 2 | 46.592 | 375400 |
| QQL06651.1;YP_005090075.1;sp\|Q2PQH4.1\|ATPB_FUCVE;CAX12455.1;ABC25543.1 | AtpB, partial (plastid) [Fucus vesiculosus] | QQL06651.1 AtpB, partial (plastid) [Fucus vesiculosus];YP_005090075.1 ATP synthase CF1 beta chain (chloroplast) [Fucus vesiculosus];sp\|Q2PQH4.1\|ATPB_FUCVE RecName: Full=ATP synthase subunit beta, chloroplastic; AltName: Full=ATP synthase F1 sector subunit | 1 | 46.294 | 101270 |
| **Ectocarpus siliculosus, Ectocarpus subulatus, Fucus, Porphyra umbilicalis, Saccharina japonica, Ulva mutabilis, Ulva prolifera** |  |  |  |  |  |
| CAB82768.1;CBJ48887.1;OSX70171.1;OSX80309.1;OSX80298.1;CBN79887.1;CBN77461.1;CBN77454.1;CBJ48892.1;CBJ33200.1;CBJ32083.1;CBJ32075.1;OSX79042.1 | histone H3, partial | CAB82768.1 histone H3, partial [Fucus serratus];CBJ48887.1 conserved unknown protein [Ectocarpus siliculosus];OSX70171.1 hypothetical protein BU14_0877s0004 [Porphyra umbilicalis];OSX80309.1 hypothetical protein BU14_0055s0031 [Porphyra umbilicalis];OSX802 | 1 | 12.302 | 9908600 |
| CBN74671.1;CBN77457.1;CBN77453.1;CBN77408.1;CBJ48885.1;CBJ32077.1;OSX71628.1;OSX71620.1;OSX81223.1;OSX77442.1;CBJ32072.1;OSX71570.1;CBJ33203.1 | histone H2A (ISS) | CBN74671.1 histone H2A (ISS) [Ectocarpus siliculosus];CBN77457.1 histone H2A isoform 2 [Ectocarpus siliculosus];CBN77453.1 histone H2A isoform 2 [Ectocarpus siliculosus];CBN77408.1 histone H2A isoform 2 [Ectocarpus siliculosus];CBJ48885.1 histone H2A (ISS) | 1 | 12.646 | 7394200 |
| WYC12533.1;WYC12531.1;WYC12530.1;CBJ30601.1;ACJ05913.1;ABB80121.1;ABB02445.1;sp\|Q39758.1\|ACT_FUCVE;CAA67388.1 | actin | WYC12533.1 actin [Saccharina japonica];WYC12531.1 actin [Saccharina japonica];WYC12530.1 actin [Saccharina japonica];CBJ30601.1 actin [Ectocarpus siliculosus];ACJ05913.1 beta-actin [Saccharina japonica];ABB80121.1 actin [Saccharina japonica];ABB02445.1 act | 3 | 41.79 | 4504100 |
| CBN77459.1;CBN77456.1;CBN77451.1;CBJ48890.1;CBJ48889.1;CBJ32081.1;CBJ32073.1;CBJ33198.1 | histone H4 | CBN77459.1 histone H4 [Ectocarpus siliculosus];CBN77456.1 histone H4 [Ectocarpus siliculosus];CBN77451.1 histone H4 [Ectocarpus siliculosus];CBJ48890.1 histone H4 [Ectocarpus siliculosus];CBJ48889.1 histone H4 [Ectocarpus siliculosus];CBJ32081.1 histone H4 | 5 | 11.411 | 4471600 |
| CBJ29997.1;OSX79791.1;CBJ31240.1 | Glyceraldehyde 3-phosphate dehydrogenase; alternative name: imm downregulated 12 | CBJ29997.1 Glyceraldehyde 3-phosphate dehydrogenase; alternative name: imm downregulated 12 [Ectocarpus siliculosus];OSX79791.1 hypothetical protein BU14_0071s0045 [Porphyra umbilicalis];CBJ31240.1 glyceraldehyde 3-phosphate dehydrogenase [Ectocarpus silic | 1 | 33.617 | 589970 |
| CBJ32894.1;CBJ32893.1;OSX75723.1;OSX74984.1;OSX75725.1 | eukaryotic translation elongation factor 1 alpha | CBJ32894.1 EEF1A1, eukaryotic translation elongation factor 1 alpha [Ectocarpus siliculosus];CBJ32893.1 EEF1A2, eukaryotic translation elongation factor 1 alpha [Ectocarpus siliculosus];OSX75723.1 hypothetical protein BU14_0224s0021 [Porphyra umbilicalis]; | 2 | 48.63 | 589850 |
| sp\|A8CEP3.1\|CALM_SACJA;CBJ33511.1;BAF80878.1 | Calmodulin, Calcium-binding protein | sp\|A8CEP3.1\|CALM_SACJA RecName: Full=Calmodulin; Short=CaM;CBJ33511.1 Calcium-binding protein [Ectocarpus siliculosus];BAF80878.1 calmodulin [Saccharina japonica] | 2 | 16.768 | 527050 |
| OSX68412.1 | hypothetical protein BU14_2849s0001 | OSX68412.1 hypothetical protein BU14_2849s0001 [Porphyra umbilicalis] | 1 | 11.545 | 498380 |
| CBJ25607.1 | flagellar associated protein | CBJ25607.1 flagellar associated protein [Ectocarpus siliculosus] | 1 | 119.74 | 213470 |
| CBN77458.1;CBN77452.1;CBN77409.1;CBN74672.1;CBJ48886.1;CBJ33202.1;CBJ32076.1 | conserved unknown protein | CBN77458.1 conserved unknown protein [Ectocarpus siliculosus];CBN77452.1 conserved unknown protein [Ectocarpus siliculosus];CBN77409.1 conserved unknown protein [Ectocarpus siliculosus];CBN74672.1 conserved unknown protein [Ectocarpus siliculosus];CBJ48886 | 1 | 12.798 | 209060 |
| AEN71225.1;ACJ05912.1;CBJ29307.1;OSX76008.1;CBJ48460.1;CBJ29295.1;CBJ32839.1;OSX71346.1;CBN79394.1 | heat shock protein 70 | AEN71225.1 heat shock protein 70 [Saccharina japonica];ACJ05912.1 heat shock protein 70 [Saccharina japonica];CBJ29307.1 Heat shock protein 70, likely pseudogene [Ectocarpus siliculosus];OSX76008.1 hypothetical protein BU14_0212s0028 [Porphyra umbilicalis] | 2 | 72.056 | 195790 |
| OSX74996.1 | hypothetical protein BU14_0258s0001 | OSX74996.1 hypothetical protein BU14_0258s0001 [Porphyra umbilicalis] | 1 | 58.183 | 139870 |
| OSX80849.1 | hypothetical protein BU14_0031s0021 | OSX80849.1 hypothetical protein BU14_0031s0021 [Porphyra umbilicalis] | 1 | 115.12 | 137700 |
| OSX69090.1 | hypothetical protein BU14_1872s0001 | OSX69090.1 hypothetical protein BU14_1872s0001 [Porphyra umbilicalis] | 1 | 89.824 | 123050 |
| CBJ32648.1 | viral A-type inclusion protein | CBJ32648.1 viral A-type inclusion protein [Ectocarpus siliculosus] | 1 | 394.76 | 105080 |
| CBN77326.1 | conserved unknown protein | CBN77326.1 conserved unknown protein [Ectocarpus siliculosus] | 1 | 50.692 | 91264 |
| CBN79446.1;CBJ30701.1;CBJ26065.1;CBN79445.1;CBJ26064.1;CON__ENSEMBL:ENSBTAP00000025008 | beta tubulin | CBN79446.1 beta tubulin [Ectocarpus siliculosus];CBJ30701.1 beta tubulin [Ectocarpus siliculosus];CBJ26065.1 beta tubulin [Ectocarpus siliculosus];CBN79445.1 beta tubulin [Ectocarpus siliculosus];CBJ26064.1 beta tubulin [Ectocarpus siliculosus]; | 1 | 50.132 | 54235 |
| CBN77378.1;CBN77373.1;CBJ28184.1;OSX79472.1 | alpha tubulin | CBN77378.1 alpha tubulin [Ectocarpus siliculosus];CBN77373.1 alpha tubulin [Ectocarpus siliculosus];CBJ28184.1 alpha tubulin [Ectocarpus siliculosus];OSX79472.1 hypothetical protein BU14_0076s0028 [Porphyra umbilicalis] | 2 | 49.994 | 49845 |
| OSX69185.1 | hypothetical protein BU14_1766s0002 | OSX69185.1 hypothetical protein BU14_1766s0002 [Porphyra umbilicalis] | 1 | 58.182 | 43043 |
| YP_003289201.1;QIE12433.1;CAV31232.1;CAT18760.1 | 30S ribosomal protein S19 | YP_003289201.1 30S ribosomal protein S19 (chloroplast) [Ectocarpus siliculosus];QIE12433.1 30S ribosomal protein S19 (plastid) [Ectocarpus siliculosus];CAV31232.1 Chloroplast 30S ribosomal protein S19 (chloroplast) [Ectocarpus siliculosus];CAT18760.1 Chlor | 1 | 10.965 | 0 |
| CBJ26903.1 | Sporulation-specific protein 11 homologue | CBJ26903.1 Sporulation-specific protein 11 homologue [Ectocarpus siliculosus] | 1 | 36.772 | 0 |
| CBJ27423.1;OSX69422.1 | proteasome subunit alpha | CBJ27423.1 proteasome subunit alpha [Ectocarpus siliculosus];OSX69422.1 hypothetical protein BU14_1526s0001 [Porphyra umbilicalis] | 1 | 25.874 | 0 |
| CBJ31855.1 | expressed unknown protein | CBJ31855.1 expressed unknown protein [Ectocarpus siliculosus] | 1 | 205.93 | 0 |
| OSX72513.1 | hypothetical protein BU14_0427s0003 | OSX72513.1 hypothetical protein BU14_0427s0003 [Porphyra umbilicalis] | 1 | 98.037 | 0 |

Fvp\_gel
| Fucus vesiculosus |  |  |  |  |  |
| --- | --- | --- | --- | --- | --- |
| Majority protein IDs | Protein names | Fasta headers | Unique peptides | Mol. weight [kDa] | iBAQ |
| YP_448618.1;AAR29312.1 | ribosomal protein S11 (mitochondrion) [Fucus vesiculosus] | YP_448618.1 ribosomal protein S11 (mitochondrion) [Fucus vesiculosus];AAR29312.1 ribosomal protein S11 (mitochondrion) [Fucus vesiculosus] | 1 | 20.553 | 858300 |
| sp\|Q39758.1\|ACT_FUCVE;CAA67388.1 | Actin; beta-actin [Fucus vesiculosus] | sp\|Q39758.1\|ACT_FUCVE RecName: Full=Actin;CAA67388.1 beta-actin [Fucus vesiculosus] | 2 | 41.889 | 641140 |
| YP_005090055.1;CAX12435.1 | ATP synthase CF1 alpha chain (chloroplast) [Fucus vesiculosus] | YP_005090055.1 ATP synthase CF1 alpha chain (chloroplast) [Fucus vesiculosus];CAX12435.1 ATP synthase CF1 alpha chain (chloroplast) [Fucus vesiculosus] | 1 | 54.095 | 285550 |
| sp\|Q39757.1\|1433_FUCVE;CAA67389.1 | 14-3-3-like protein [Fucus vesiculosus] | sp\|Q39757.1\|1433_FUCVE RecName: Full=14-3-3-like protein;CAA67389.1 14-3-3 [Fucus vesiculosus] | 2 | 28.169 | 142890 |
| YP_005090075.1;sp\|Q2PQH4.1\|ATPB_FUCVE;CAX12455.1;ABC25543.1;QQL06651.1 | ATP synthase CF1 beta chain (chloroplast) [Fucus vesiculosus] | YP_005090075.1 ATP synthase CF1 beta chain (chloroplast) [Fucus vesiculosus];sp\|Q2PQH4.1\|ATPB_FUCVE RecName: Full=ATP synthase subunit beta, chloroplastic; AltName: Full=ATP synthase F1 sector subunit beta; AltName: Full=F-ATPase subunit beta;CAX12455.1 AT | 2 | 51.572 | 99756 |
| YP_005090045.1;CAX12425.1 | DNA-directed RNA polymerase beta chain (chloroplast) [Fucus vesiculosus] | YP_005090045.1 DNA-directed RNA polymerase beta chain (chloroplast) [Fucus vesiculosus];CAX12425.1 DNA-directed RNA polymerase beta chain (chloroplast) [Fucus vesiculosus] | 1 | 127.5 | 33982 |
| YP_005090125.1;CAX12511.1 | cell division protein FtsH-like protein (chloroplast) [Fucus vesiculosus] | YP_005090125.1 cell division protein FtsH-like protein (chloroplast) [Fucus vesiculosus];CAX12511.1 cell division protein FtsH-like protein (chloroplast) [Fucus vesiculosus] | 2 | 69.652 | 22247 |
| **Ectocarpus siliculosus, Ectocarpus subulatus, Fucus, Porphyra umbilicalis, Saccharina japonica, Ulva mutabilis, Ulva prolifera** |  |  |  |  |  |
| OSX79327.1 | hypothetical protein BU14_0081s0007 | OSX79327.1 hypothetical protein BU14_0081s0007 [Porphyra umbilicalis] | 1 | 26.907 | 52209000 |
| QID90270.1 | sulfotransferase | QID90270.1 sulfotransferase [Saccharina japonica] | 1 | 48.359 | 10542000 |
| OSX76083.1 | hypothetical protein BU14_0207s0002 | OSX76083.1 hypothetical protein BU14_0207s0002 [Porphyra umbilicalis] | 1 | 56.643 | 4728200 |
| AUO68231.1;AUO68228.1;AUO68227.1;QED55811.1;AUO68237.1;AUO68236.1;AUO68234.1;AUO68233.1;AUO68232.1;AUO68230.1;AUO68229.1;AUO68226.1;AUO68225.1;AUO68224.1 | Tic20-like protein | AUO68231.1 Tic20-like protein [Saccharina japonica];AUO68228.1 Tic20-like protein [Saccharina japonica];AUO68227.1 Tic20-like protein [Saccharina japonica];QED55811.1 chloroplast Tic20-14 protein precursor [Saccharina japonica];AUO68237.1 Tic20-like protei | 1 | 43.818 | 1827100 |
| WYC12533.1;WYC12531.1;WYC12530.1;sp\|Q39758.1\|ACT_FUCVE;CBJ30601.1;CAA67388.1;ACJ05913.1;ABB80121.1;ABB02445.1 | actin | WYC12533.1 actin [Saccharina japonica];WYC12531.1 actin [Saccharina japonica];WYC12530.1 actin [Saccharina japonica];sp\|Q39758.1\|ACT_FUCVE RecName: Full=Actin;CBJ30601.1 actin [Ectocarpus siliculosus];CAA67388.1 beta-actin [Fucus vesiculosus];ACJ05913.1 be | 1 | 41.79 | 469830 |
| OSX72195.1;BAS06569.1;OSX79458.1 | hypothetical protein BU14_0459s0016 | OSX72195.1 hypothetical protein BU14_0459s0016 [Porphyra umbilicalis];BAS06569.1 xanthoxin dehydrogenase [Porphyra umbilicalis];OSX79458.1 hypothetical protein BU14_0076s0014 [Porphyra umbilicalis] | 1 | 33.669 | 411350 |
| OSX74742.1 | hypothetical protein BU14_0269s0022 | OSX74742.1 hypothetical protein BU14_0269s0022 [Porphyra umbilicalis] | 1 | 19.134 | 261660 |
| OSX74489.1 | hypothetical protein<br>BU14_0286s0002 | OSX74489.1 hypothetical protein BU14_0286s0002, partial [Porphyra umbilicalis] | 1 | 63.167 | 180850 |
| CBJ32648.1 | viral A-type inclusion protein | CBJ32648.1 viral A-type inclusion protein [Ectocarpus siliculosus] | 1 | 394.76 | 179640 |
| CBJ48979.1 | non-specific serine/threonine<br>protein kinase, EC:2.7.11.1 | CBJ48979.1 Protein kinase domain containing protein [Ectocarpus siliculosus] | 1 | 91.137 | 140750 |
| OSX71107.1;OSX78504.1 | hypothetical protein<br>BU14_0592s0002 | OSX71107.1 hypothetical protein BU14_0592s0002 [Porphyra umbilicalis];OSX78504.1 hypothetical protein BU14_0107s0012 [Porphyra umbilicalis] | 1 | 43.324 | 112930 |
| CBJ32676.1 | glucose regulated protein<br>/BiP | CBJ32676.1 glucose regulated protein /BiP [Ectocarpus siliculosus] | 1 | 31.681 | 78105 |
| OSX68984.1 | hypothetical protein<br>BU14_2001s0001 | OSX68984.1 hypothetical protein BU14_2001s0001 [Porphyra umbilicalis] | 1 | 75.466 | 33700 |
| OSX69185.1 | hypothetical protein<br>BU14_1766s0002 | OSX69185.1 hypothetical protein BU14_1766s0002 [Porphyra umbilicalis] | 1 | 58.182 | 0 |
| OSX78582.1 | hypothetical protein<br>BU14_0106s0047 | OSX78582.1 hypothetical protein BU14_0106s0047 [Porphyra umbilicalis] | 1 | 103.64 | 0 |

## Notes

### Competing Interest Statement

The authors have declared no competing interest.

